# Optimizing Avian Influenza Viral RNA Enrichment from Water: A Comparison of Concentration Methods for Low-Volume Samples

**DOI:** 10.64898/2026.11.13.766378

**Authors:** Joaquín Neumann Heise, Chiara Holzer, Claudia Stange, Andreas Tiehm, Albert Perlas, Lara Urban, Timm Harder, Lutz Breuer, Sascha Knauf, Anja Globig

**Author notes:** **Correspondence**: Anja Globig.

## Abstract

Feed-baited water bins may provide a complementary approach for avian influenza virus (AIV) surveillance in wild waterbird populations. In a previous study, water from these bins was sampled using swabs after exposure to visiting birds, but the low sample volume limited the detection and subsequent characterization of AIV RNA (F. I. Rondi, J. Neumann-Heise, T. Harder, V. Allendorf, K. Depner, T. Homeier-Bachmann, L. Breuer, S. Knauf, S. Dreyer, and A. Globig, One Health 23:101548, 2026, https://doi.org/10.1016/j.onehlt.2026.101548). Here, we compared five sample-processing methods for improving AIV RNA detection in small (40-mL) water samples (tap and brackish water) spiked with an AIV of the H4N6 subtype under simulated field conditions: swab sampling, polyethylene glycol precipitation, activated carbon capture, combined activated carbon capture and polyethylene glycol precipitation (AC/PEG), and electronegative membrane adsorption (EM). AC/PEG and EM showed the strongest overall performance based on RT-qPCR results, although their relative performance varied across experimental conditions. The qPCR-derived viral signal was generally higher in tap water than in brackish water, whereas wheat presence was associated with a lower signal, particularly for swab sampling. Subsequent testing was conducted to ascertain the most effective storage conditions with regard to temperature and duration. During storage, AIV RNA detection remained relatively stable at −20 °C for up to four weeks, with only minor increases in mean Ct values in both water matrices. Greater increases were observed at room temperature and 4 °C, with significantly higher Ct values at both temperatures than at −20 °C in both water matrices at Weeks 2 and 4 (*p* < 0.05). These findings support the potential of AC/PEG and EM to facilitate the recovery, detection and subsequent characterization of AIV from low-volume water samples by improving viral RNA detection using RT-qPCR. Given its overall performance and lower cost compared with EM, AC/PEG may represent a practical option for field-based surveillance. Regarding storage, −20 °C provided the best conditions for maintaining AIV RNA detection, with relatively stable results for up to four weeks.

**Importance:** Environmental surveillance can complement conventional approaches for detecting avian influenza virus (AIV) circulation in wild waterbirds. Feed-baited water bins offer a non-invasive sampling strategy, but processing low-volume water samples may require viral enrichment to improve AIV RNA detection. Here, we show that combined activated carbon capture and polyethylene glycol precipitation (AC/PEG) is a promising, cost-effective method for improving AIV RNA detection in low-volume water samples. These findings highlight the potential of viral enrichment methods to support the use of simple, practical and non-invasive approaches for AIV surveillance.

## 1. Introduction

Among wild waterbirds, the orders Anseriformes (ducks, geese, and swans) and Charadriiformes (gulls, terns, and waders) form a natural reservoir system for avian influenza viruses (AIV), including viruses of the H1–H16 and N1–N9 subtypes (Munster *et al*., 2007), as well as the recently described H19 subtype (Fereidouni *et al*., 2023).

Surveillance of AIV in wild bird populations provides important information on virus circulation and epidemiology. Although passive surveillance, interpreted here as the examination of sick and dead birds, is particularly valuable for detecting high pathogenicity avian influenza (HPAI)-associated mortality, active surveillance of apparently healthy birds provides complementary information on AIV circulation that may otherwise go undetected. This is particularly relevant for H5 and H7 viruses, as low-pathogenicity viruses of these subtypes may eventually acquire a high pathogenicity phenotype through spontaneous mutation after introduction into poultry populations (WOAH, 2023), although other pathways of emergence cannot be excluded.

Obtaining samples from living wild birds, however, requires capture, handling, and individual sampling. Not only has this the potential to cause considerable welfare issues as well as disturbance of ecosystems, but it can also be time- and resource-intensive, particularly when frequent sampling is required. These practical constraints have encouraged the development of complementary approaches that do not depend on direct sampling of individual birds.

AIV infection in waterfowl can result in viral shedding through both oral and cloacal routes (Hénaux and Samuel, 2011), providing opportunities for viral material to enter the environment and contaminate different matrices, including surface water, sediment, faeces (Pannwitz *et al*., 2009; Ahrens *et al*., 2023), and feathers (Azeem *et al*., 2023).

Among these matrices, surface water is particularly relevant because viral material shed by multiple wild waterbirds can accumulate in the same aquatic environment. Accordingly, AIV RNA has been detected in water bodies frequented by wild waterbirds, although viral concentrations can be low, limiting the sensitivity of molecular detection and potentially constraining subsequent virus characterization (Germeraad *et al*., 2020; Ahrens *et al*., 2023; Hubbard *et al*., 2023).

Several approaches have been investigated to concentrate viruses from environmental water, including filtration, ultrafiltration, precipitation, adsorption, and combinations of these methods (Cashdollar and Wymer, 2013). The amount of viral material available for detection after processing depends in part on the volume of water processed, making efficient virus enrichment particularly important for low-volume samples (Coombe *et al*., 2021).

Feed-baited water bins as a decoy for wild waterbirds have been proposed as a complementary approach for environmental surveillance of AIV (Rondi *et al*., 2026). In this setup, shallow water bins containing wheat covered with water as a feed source are used to attract wild waterbirds. Following their visits, during which they may leave AIV in the water, swabs are used to collect samples from the water in the bins, which are then tested for AIV. However, AIV RNA was detected in only 13 of 222 water samples (5.9%), and nine positive samples could not be subtyped due to low viral loads (Rondi *et al*., 2026). Although some samples may genuinely lack AIV RNA, improved sample-processing methods, particularly those involving viral RNA concentration, may help enable reliable subtyping and reduce the risk of false-negative results in samples with low viral loads.

To address this need, the present study compared five sample-processing methods using low-volume (40-mL) water samples under controlled laboratory conditions designed to simulate the feed-baited water-bin approach. These comprised direct swab sampling, which does not involve an enrichment step, and four enrichment methods. The aim was to identify an effective, practical, and cost-effective method for subsequent field application. In addition, we assessed the effects of water matrix and wheat presence on the qPCR-derived viral signal and investigated how storage temperature and duration affected AIV RNA detection.

We hypothesized that AIV RNA detection would differ among the five sample-processing methods, with the four enrichment methods expected to yield higher qPCR-derived viral signals than direct swab sampling. We further hypothesized that the magnitude of this difference would be influenced by water matrix and wheat presence.

## 2. Materials and Methods

### 2.1. Experimental design

The study was conducted in two sequential experiments. Experiment 1 compared five sample-processing methods: swab sampling (S), polyethylene glycol precipitation (PEG), activated carbon capture (AC), combined activated carbon capture and PEG (AC/PEG), and electronegative membrane adsorption (EM). Unlike S, which involved direct sampling without enrichment, PEG, AC, AC/PEG, and EM involved enrichment of the viral material. The methods were evaluated using two water matrices (tap and brackish water), each with and without wheat. When wheat was included, 5 g of wheat grains were added to each 40-mL water sample, reflecting its use as bait to attract waterbirds to the water bins (Rondi *et al*., 2026).

Based on the results of Experiment 1, AC/PEG was selected for Experiment 2 based on its overall performance in improving AIV RNA detection and its lower cost compared with EM. Experiment 2 evaluated the effects of storage temperature and duration on AIV RNA detection using both water matrices without wheat.

### 2.2. Water matrices

Two water matrices were used: non-chlorinated tap water and brackish surface water collected from the Greifswalder Bodden, a coastal bay in the southwestern Baltic Sea, northeastern Germany. Before use, brackish water samples were screened for AIV RNA using one or more of the methods evaluated in this study (Supplementary Table S10). Methods that repeatedly yielded no detectable AIV RNA while others detected AIV were discontinued for subsequent samples to avoid unnecessary processing. Only samples with no detectable AIV RNA across the applied methods were included in the experiments. Prior to the experiments, the tap water had an average salinity of 0.92 ‰ and a pH of 7.1, whereas the brackish water had an average salinity of 7.3 ‰ and an average pH of 6.9. Sample preparation and sample-processing procedures were conducted at room temperature, with initial water temperatures ranging from 21 to 24 °C.

### 2.3. Virus

Influenza A virus A/rhea/Germany/AI06884/2023 (H4N6) had been isolated from an oropharyngeal swab sample of a Greater Rhea (*Rhea americana*) in a zoo. The virus originated from the virus repository at the National Reference Laboratory for Avian Influenza, Friedrich-Loeffler-Institut, Germany, and was amplified for up to 72 h in the allantoic cavity of 9-day-old embryonated chicken eggs. Allantoic fluid was harvested and virus infectivity titrated on Leghorn Male Hepatoma (LMH) cell culture in microtitre plates as previously described (WHO, 2011). Viral cytopathic effects developing in serum-free cultures to which TPCK-trypsin had been added at a concentration of 2 µg/mL were read by light microscopy after 72–96 h of incubation. Viral suspensions had titres of 106 to 106.5 TCID50/mL as calculated by the Kaerber method (Kaerber, 1931).

### 2.4. Sample preparation

Sterile 50 mL DNase-/RNase-free polypropylene tubes (Sarstedt, Nümbrecht, Germany) were filled with 40 mL of the corresponding water matrix. Based on an initial virus stock of 10⁶ TCID₅₀/mL, 4, 40, and 400 µL of it were added to obtain final virus concentrations of arithmetically 10², 10³, and 10⁴ TCID₅₀/mL, respectively.

In Experiment 1, three independent replicates were prepared for each combination of experimental condition, viral concentration, and sample-processing method, as well as for non-enriched samples. The non-enriched controls bypassed the sample-processing procedures and were subjected directly to RNA extraction.

A negative control consisting of specific pathogen free allantoic fluid was included in each experiment using spiking volumes equivalent to those of the virus-spiked samples.

For Experiment 2, samples were spiked to a final concentration of 10³ TCID₅₀/mL and stored at room temperature (21–24 °C), 4 °C, or −20 °C. AIV RNA detection was assessed immediately after spiking (Day 0) and after 2 and 4 weeks of storage. Three independent replicates were prepared for each combination of water matrix, storage temperature, and storage duration. Specific pathogen free allantoic fluid served as the negative control throughout the experiment.

### 2.5. Sample-processing methods

*Swab sampling* was performed essentially as described by Rondi *et al*. (2026) using ∑-VIROCULT® swabs (Check Diagnostics GmbH, Bad Oldesloe, Germany). Each swab was immersed in the corresponding water sample for 5 s and subsequently transferred into its original collection tube containing 1 mL of transport medium (balanced salt solution with buffering agents and antimicrobial supplements). The collection tubes were incubated at 4 °C for up to 15 min, briefly agitated manually, and then processed for RNA extraction.

PEG was performed according to Holzer *et al*. (2025) and Ho *et al*. (2022), with minor modifications. Briefly, spiked water samples were first centrifuged at 5000 × g for 30 min at 4 °C to remove suspended solids. Although this centrifugation step was primarily intended to remove suspended solids from the brackish water, it was also applied to tap water samples to maintain an identical sample processing protocol for both water matrices. The supernatant was then transferred to a new tube containing PEG 8000 (10% w/v; ROTIPURAN® Ph. Eur., Carl Roth, Karlsruhe, Germany) and NaCl (≥99.5%, p.a., ACS, ISO; Carl Roth, Karlsruhe, Germany). For tap water samples, NaCl was added to reach a final concentration of 2% (w/v), whereas for brackish water samples, the native salinity was measured using a multiparameter probe (Extech EC600; Extech Instruments, Nashua, NH, USA), and the amount of NaCl added was adjusted accordingly to reach the same final concentration of 2% (w/v). The mixture was incubated on ice with shaking at 400 rpm until complete dissolution. Precipitation was achieved by centrifugation at 12,000 × g for 1 h 45 min at 4 °C. The supernatant was discarded, and the resulting pellet resuspended in 300 µL of nuclease-free water (DNase/RNase-free; Invitrogen, Carlsbad, CA, USA) for subsequent RNA extraction.

The AC method also followed Holzer *et al*. (2025) with minor modifications. Briefly, 1 g of granular activated carbon (Filtrasorb 300-M, Calgon Carbon Corporation, Pittsburgh, PA, USA) was enclosed in nylon mesh bags (5 × 6 cm), which were sealed prior to use and placed into the corresponding spiked water samples. The samples were incubated overnight (16 h) at room temperature under continuous agitation at 600 rpm using an orbital shaker to allow virus adsorption. The carbon was then recovered, and viruses were eluted using an alkaline beef extract elution buffer (EB4/PBS, pH ∼11) based on the formulation described by Holzer et al. (2025). The eluate (7 mL) was subsequently processed for RNA extraction.

For the combined AC/PEG approach, AC capture and elution were performed as described above. The resulting eluate was subsequently subjected to PEG following the same protocol described for PEG alone. The amounts of PEG 8000 and NaCl were adjusted proportionally to the eluate volume (7 mL) to achieve the final concentrations indicated above.

For *EM-based adsorption*, one membrane filter (EZ-PAK® cellulose membrane filters, Merck, Darmstadt, Germany, 47 mm, pore size 0.45 µm) was placed overnight (i.e. 16 h) into each corresponding spiked water sample and incubated under the same conditions as for AC capture.

### 2.6. RNA extraction

For samples processed using S, PEG, AC, or AC/PEG, the resulting sample fractions were subjected to manual viral RNA extraction using the QIAamp RNA Mini Kit (Qiagen, Hilden, Germany). For samples processed using the EM method, the electronegative membrane itself was transferred directly to the RNeasy PowerWater Kit (Qiagen, Hilden, Germany) for simultaneous elution of viral material and RNA extraction. RNA extraction was performed in batches corresponding to each experiment. Extracted RNA from each batch was stored at −80 °C until all samples had been processed.

A heterologous external transcription and amplification control (IC2 RNA) was added during RNA extraction for all samples, regardless of the extraction method used, as previously described (Hoffmann *et al*., 2006), also allowing the detection of potential inhibitory effects. In addition, a negative extraction control, consisting of the transport medium supplied with VIROCULT® swabs described above, was included in each batch.

PCR inhibitory effects were observed in preliminary experiments (TZW, unpublished data) with samples processed using activated carbon. Therefore, for samples processed using AC or AC/PEG, the extracted RNA was further purified using the OneStep PCR Inhibitor Removal Kit (Zymo Research, Irvine, CA, USA) to reduce PCR inhibition. This observation is consistent with previous reports describing the adsorption of PCR-inhibitory compounds by activated carbon (Abolmaaty *et al*., 2007; Schrader *et al*., 2012).

### 2.7. Real-time reverse transcription–PCR (RT-qPCR)

Detection of influenza A virus was performed using a generic influenza A virus RT-qPCR assay targeting the matrix gene (Spackman *et al*., 2002), modified as previously described (Hassan *et al*., 2022), and conducted using the AgPath-ID™ One-Step RT-qPCR Reagents kit (Applied Biosystems™, Thermo Fisher Scientific, Foster City, CA, USA) on a CFX Opus 96 Real-Time PCR System (Bio-Rad Laboratories, Hercules, CA, USA). A no-template control was included in every RT-qPCR run. Threshold cycle (Ct) values were used as the primary measure of viral detection and served as the basis for subsequent statistical analyses.

### 2.8. Quantification and normalization of viral signal

#### 2.8.1. Conversion of Ct values to TCID₅₀-equivalent (TCID₅₀-eq) genome quantities

A standard curve was generated using four independent series of 1:10 dilutions of RNA extracted from the H4N6 virus stock, covering nominal concentrations of 10⁶ to 10⁻¹ TCID₅₀/mL. Each series was initiated from a separate aliquot of the same RNA preparation corresponding to 10⁶ TCID₅₀/mL, resulting in four replicates per concentration (Supplementary Table S7). The relationship between Ct values and the logarithm of the corresponding TCID₅₀ concentration was assessed by linear regression using the CFX Opus 96 system software, which automatically calculated the slope (−3.474), intercept (29.119), amplification efficiency (97.6%), and R² (0.991).

Ct values from the experimental samples were subsequently converted to TCID₅₀-eq using the inverse standard-curve equation for quantitative comparison of the qPCR-derived viral signal among sample-processing methods:

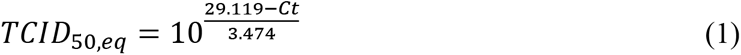

#### 2.8.2. Relative qPCR-derived viral signal

For each sample-processing method, viral concentration, and experimental condition, the mean TCID₅₀-eq of the three independent replicates was divided by the corresponding mean TCID₅₀-eq of the three non-enriched controls to calculate fold-change (*FC*) in qPCR-derived viral signal relative to the non-enriched control, as follows:

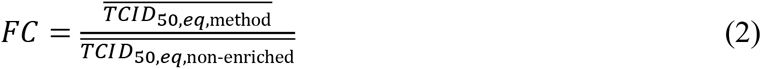

#### 2.8.3. Equivalent sample volume-based normalization of the qPCR-derived viral signal

The primary assessment of sample-processing method performance was based on RT-qPCR Ct values, as these measurements reflect the analytical performance of each method as applied under practical conditions. As a complementary analysis, an equivalent sample volume (ESV)-based normalization was performed to account for differences among methods in the volume of original sample represented in the RT-qPCR reaction, resulting from differences in the volume of material recovered and the fraction subjected to RNA extraction (Borchardt *et al*., 2021; Crank *et al*., 2023).

For methods yielding a liquid concentrate, ESV was calculated from the initial sample volume *V_intial_*, the recovered sample volume *V_recovered_*, the volume subjected to RNA extraction *V_extraction_*, the RNA elution volume *V_elution_*, and the volume of RNA used in the RT-qPCR reaction *V_PCR_*:

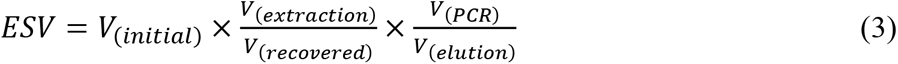

Because reverse transcription and qPCR were performed in a single reaction, the additional volume correction for transfer of cDNA to a separate PCR reaction described by Crank *et al*. (2023) was not applicable.

For EM, which generated no intermediate liquid concentrate, no recovered-volume term was included, and the *ESV_EM_* was calculated as follows:

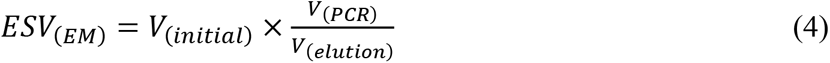

Details of the volumetric parameters used in ESV calculations for each method are listed in Table 1.

**Table 1.** Volumetric parameters and Equivalent Sample Volumes (ESV) used for normalization. ESV represents the volume of the original 40-mL sample represented in one RT-qPCR reaction. For EM, RNA was extracted directly from the processed membrane; therefore, no recovered liquid volume was used in the ESV calculation. Abbreviations: S, *swab sampling*; PEG, *polyethylene glycol precipitation*; AC, *activated carbon capture*; AC/PEG, *activated carbon capture* combined with PEG; EM, *electronegative membrane-based adsorption*.

| Sample-processing method | Initial sample volume | Volume processed | Final recovered volume | Volume used for RNA extraction | RNA elution volume | RNA used for RT-qPCR | Equivalent Sample Volume (ESV) |
|---|---|---|---|---|---|---|---|
| Non-enriched | 40 mL | No enrichment; direct aliquot | — | 140 µL | 50 µL | 5 µL | 0.014 mL |
| S | 40 mL | Swab immersed in the entire 40 mL sample | 1,000 µL transport medium | 140 µL | 50 µL | 5 µL | 0.560 mL |
| PEG | 40 mL | 40 mL | 300 µL | 140 µL | 50 µL | 5 µL | 1.867 mL |
| AC | 40 mL | 40 mL; AC nylon bag incubated in sample | 7,000 µL | 140 µL | 50 µL | 5 µL | 0.080 mL |
| AC/PEG | 40 mL | 40 mL; AC nylon bag incubated in sample | 300 µL | 140 µL | 50 µL | 5 µL | 1.867 mL |
| EM | 40 mL | 40 mL; membrane incubated in sample | — | Material on membrane processed directly | 100 µL | 5 µL | 2.000 mL |

ESV-normalized qPCR-derived viral signal was expressed relative to the corresponding non-enriched control after normalization of both signals to their respective ESV values (equation 5). This metric represents recovery relative to the corresponding non-enriched reference rather than absolute recovery relative to the nominal amount of virus spiked.

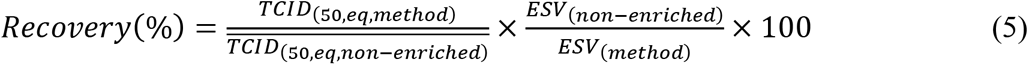

### 2.9. Effect of storage conditions on AIV RNA detection

The sample-processing method selected based on the results of Experiment 1 was used to evaluate the effects of storage conditions relevant to field applications on AIV RNA detection. Water samples were prepared using tap water and brackish water without wheat and spiked at 10³ TCID₅₀/mL. Samples were stored at room temperature (21–24 °C), 4 °C, or −20 °C and analyzed at Day 0 and after 2 and 4 weeks of storage. Three independent replicates were prepared for each combination of water matrix and storage condition.

At each sampling time, viral RNA was extracted and quantified using the same RT-qPCR procedure described above. AIV RNA detection was assessed based on the resulting Ct values.

### 2.10. Statistical analysis

#### 2.10.1. Software

Statistical analyses were performed using R version 4.3.1 (R Core Team, 2023). Data import and processing were performed using *readxl* v1.4.3 (Wickham and Bryan, 2023) and *dplyr* v1.1.4 (Wickham *et al*., 2023), respectively. Statistical models were fitted using R, with Type III ANOVA performed using car v3.1-3 (Fox and Weisberg, 2019) and estimated marginal means calculated using emmeans v2.0.1 (Lenth and Piaskowski, 2025). Data visualization and figure assembly were performed using ggplot2 v3.5.1 (Wickham, 2016), scales v1.3.0 (Wickham *et al*., 2023), and patchwork v1.2.0 (Pedersen, 2024).

#### 2.10.2. Comparison of sample-processing methods based on Ct values

For each water matrix and wheat condition, the relationship between viral concentration and RT-qPCR cycle threshold (Ct) was evaluated separately for each sample-processing method against the corresponding non-enriched control. Individual replicate Ct values were used for the analyses, with viral concentration expressed as log_10_-transformed TCID_50_/mL. For each sample-processing method, a separate Analysis of Covariance (ANCOVA) was fitted including the sample-processing method and corresponding non-enriched control as the two levels of the categorical factor *Method*. The model was specified as:

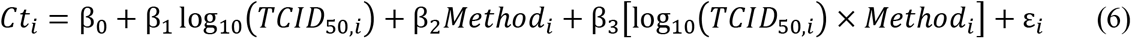

where Ct*_i_* is the Ct value of observation *i*, with *i* denoting each individual experimental replicate; log_10_(TCID_50_,_i_) represents the log-transformed viral concentration for observation *i*; and *Method* is a categorical variable with the non-enriched control as the reference level and the respective sample-processing method as the second level. β_0_ represents the intercept of the non-enriched control, β_1_ its regression slope, β_2_ the difference in intercepts between the respective sample-processing method and the non-enriched control, and β_3_ the difference in regression slopes between the two methods. The term ε_i_ represents the residual error for observation *i*.

The interaction term was tested to determine whether the relationship between viral concentration and Ct differed between the sample-processing method and the corresponding non-enriched control. When the interaction was significant (*p* < 0.05), estimated differences in Ct between the sample-processing method and the non-enriched control were evaluated separately at each of the three pre-specified viral concentrations (10², 10³, and 10⁴ TCID₅₀/mL) using estimated marginal means from the final model. These concentration-specific comparisons were defined *a priori* as planned comparisons to characterize the interaction across the experimental concentration range. *P*-values for the concentration-specific comparisons were adjusted for multiple comparisons using the Holm method within each method × experimental condition. When the interaction was not significant (*p* ≥ 0.05), a reduced ANCOVA model without the interaction term was fitted:

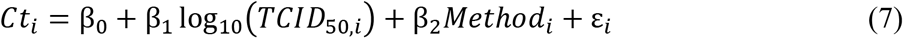

In this reduced model, β₂ represents the adjusted difference in Ct between the sample-processing method and the non-enriched control, assuming parallel regression slopes. Model assumptions were assessed by inspection of residual plots, including linearity, homoscedasticity, and approximate normality of the residuals. The model fit was assessed using R² and adjusted R² values obtained from the final ANCOVA models. A significance level of α = 0.05 was used throughout the analyses. Regression lines shown in Figure 1 were generated from the final model selected for each sample-processing method comparison.

**Figure 1.**
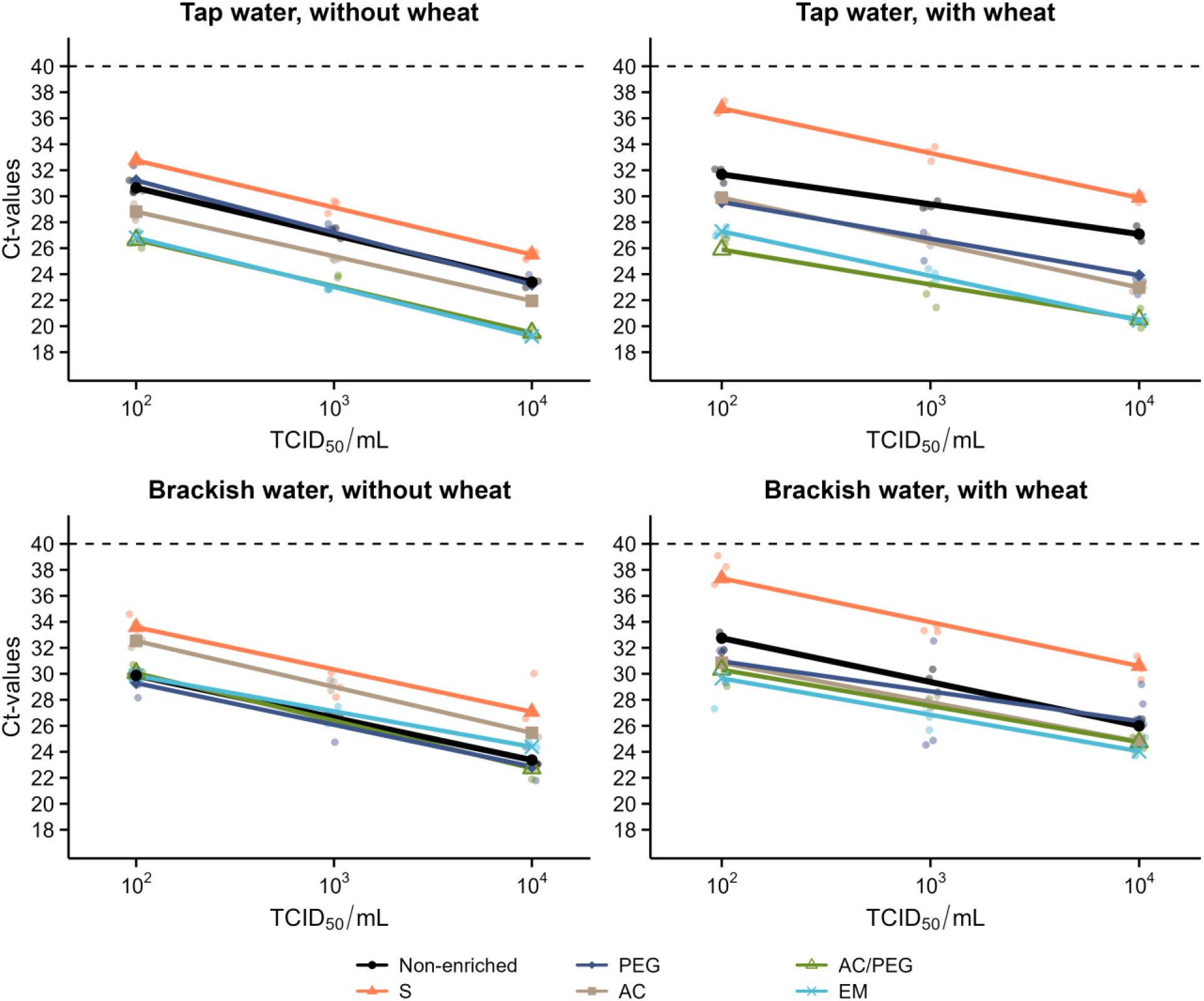
Relationship between spiked H4N6 concentration and RT-qPCR Ct values across sample-processing methods, water matrices, and wheat conditions. Individual points represent independent technical replicates (n = 3 per concentration and method), and lines represent fitted regression lines. Regression lines for sample-processing methods were generated from the final ANCOVA models, whereas the non-enriched control line was fitted separately to the corresponding control data. The dashed horizontal line indicates the RT-qPCR cut-off value (Ct = 40). Abbreviations: S, *swab sampling*; PEG, *polyethylene glycol precipitation*; AC, *activated carbon capture*; AC/PEG, *activated carbon capture* combined with PEG; EM, *electronegative membrane-based adsorption*.

#### 2.10.3. Analysis of AIV RNA detection under different storage conditions

For Experiment 2, the effects of storage temperature, storage duration, and water matrix on the qPCR-derived viral signal were evaluated using a factorial linear model, with Ct values as the response variable. Overall effects and interactions in the final model were assessed using Type III ANOVA. Day 0 samples were excluded from the factorial analysis because they were analyzed immediately after sample preparation and were not subjected to a storage temperature; they were therefore used as a descriptive baseline for comparison with the stored samples. Water matrix, storage temperature, and storage duration were included as explanatory factors, together with their two-way interactions (Water × Temperature, Water × Week, and Temperature × Week). The three-way interaction (Water × Temperature × Week) was initially included in the model but was removed from the final model because its exclusion did not result in a significant loss of model fit (F = 2.55, *p* = 0.099). Model assumptions were assessed by inspection of residual plots, including homoscedasticity and approximate normality of the residuals. Estimated marginal means were calculated, and pairwise comparisons were performed to assess changes between Week 2 and Week 4 within each water matrix and storage temperature. A significance level of α = 0.05 was used throughout the analyses.

## 3. Results

### 3.1. Comparison of sample-processing methods based on Ct values

The qPCR-derived viral signal generally increased with increasing spiked H4N6 concentration across the experimental conditions. However, the magnitude of the differences in qPCR-derived viral signal between methods and the corresponding non-enriched controls varied depending on the water matrix, wheat condition, and spiked H4N6 concentration (Figure 1; Supplementary Figure S1). The distribution of individual replicate-level Ct values across the experimental conditions is shown in Supplementary Figure S2, and the corresponding raw Ct values are provided in Supplementary Table S8.

In tap water without wheat, none of the methods showed a significant H4N6 concentration × method interaction (all *p* ≥ 0.102), indicating parallel relationships between Ct and spiked H4N6 concentration. AC/PEG and EM yielded substantially higher qPCR-derived viral signals than the non-enriched control, followed by AC. PEG did not differ significantly from the control, whereas S yielded a significantly lower qPCR-derived viral signal (Supplementary Table S1).

In tap water with wheat, significant H4N6 concentration × method interactions were detected for EM, AC, and S (all *p* < 0.001), indicating concentration-dependent differences relative to the non-enriched control. The qPCR-derived viral signal increased with increasing spiked H4N6 concentration for EM and AC, whereas S consistently yielded a lower signal than the control. No significant interaction was detected for AC/PEG or PEG, and both methods yielded higher qPCR-derived viral signals than the control (Supplementary Tables S1 and S2).

In brackish water without wheat, a significant H4N6 concentration × method interaction was detected only for EM (*p* = 0.005). Its signal did not differ significantly from the non-enriched control at 10² TCID₅₀/mL, but yielded a lower qPCR-derived viral signal at 10³ and 10⁴ TCID₅₀/mL (Supplementary Table S2). AC/PEG and PEG did not differ significantly from the control, whereas AC and S yielded significantly lower qPCR-derived viral signals (Supplementary Table S1).

In brackish water with wheat, none of the sample-processing methods showed a significant H4N6 concentration × method interaction (all *p* ≥ 0.076). AC/PEG, EM, and AC yielded significantly higher qPCR-derived viral signals than the non-enriched control. PEG did not differ significantly from the control, whereas S yielded a significantly lower qPCR-derived viral signal (Supplementary Table S1).

The goodness of fit of the final models varied among sample-processing methods and experimental conditions, with R² values predominantly ranging from 0.89 to 0.99 (mean R² = 0.94); one model showed a substantially lower R² of 0.50 (Supplementary Table S1).

The corresponding relative qPCR-derived viral signals, expressed as fold changes relative to the corresponding non-enriched controls, are summarized in Table 2. In tap water, AC/PEG and EM generally produced the greatest increases in qPCR-derived viral signal, particularly in the presence of wheat. Without wheat, AC/PEG increased the signal by 12–17-fold across the three spiked H4N6 concentrations, while EM increased it by 11–18-fold. With wheat, the increases were substantially greater, reaching 110-fold for AC/PEG at 10³ TCID₅₀/mL and 83-fold for EM at 10⁴ TCID₅₀/mL. AC also increased the qPCR-derived signal in tap water, although to a lesser extent, whereas S consistently produced signals below the non-enriched reference.

**Table 2.** Values represent fold-change (FC) relative to the corresponding non-enriched control at the same viral concentration and under the same experimental condition. An FC of 1 indicates a qPCR-derived viral signal equivalent to that of the non-enriched control, whereas values >1 indicate a gain in signal (i.e., enrichment) and values <1 indicate a loss of signal. Bold values indicate the highest fold-change (FC) relative to the corresponding non-enriched control for each experimental condition and viral concentration. Abbreviations: S, *swab sampling*; PEG, *polyethylene glycol precipitation*; AC, *activated carbon capture*; AC/PEG, *activated carbon capture* combined with PEG; EM, *electronegative membrane-based adsorption*.

| **Experimental condition** | **Method** | **10<sup>2</sup> FC** | **10<sup>3</sup> FC** | **10<sup>4</sup> FC** |
|---|---|---|---|---|
| Tap water, without wheat | S | 0.26 | 0.26 | 0.22 |
|  | PEG | 0.70 | 0.77 | 1.59 |
|  | AC | 3.77 | 3.90 | 1.96 |
|  | AC/PEG | **17.46** | 11.64 | 12.36 |
|  | EM | 11.03 | **18.13** | **13.85** |
| Tap water, with wheat | S | 0.03 | 0.07 | 0.16 |
|  | PEG | 3.26 | 7.30 | 13.79 |
|  | AC | 3.31 | 6.32 | 15.54 |
|  | AC/PEG | **26.88** | **110.00** | 80.09 |
|  | EM | 18.68 | 35.04 | **82.53** |
| Brackish water, without wheat | S | 0.07 | 0.25 | 0.06 |
|  | PEG | **1.99** | **2.14** | 1.18 |
|  | AC | 0.19 | 0.21 | 0.23 |
|  | AC/PEG | 0.90 | 1.18 | **1.56** |
|  | EM | 1.01 | 1.05 | 0.38 |
| Brackish water, with wheat | S | 0.02 | 0.07 | 0.07 |
|  | PEG | 2.85 | **14.64** | 0.58 |
|  | AC | 2.25 | 2.55 | 3.31 |
|  | AC/PEG | 4.34 | 3.00 | 2.91 |
|  | EM | **9.34** | 7.34 | **3.98** |

In brackish water, the relative increases were generally smaller and more variable. Without wheat, most methods produced signals close to or below the non-enriched reference, with the exception of PEG, which produced approximately twofold higher signals at 10² and 10³ TCID₅₀/mL. In the presence of wheat, AC/PEG, AC, and EM generally produced higher signals than the non-enriched control, whereas PEG showed marked variation across spiked H4N6 concentrations (range = 0.58–14.64-fold).

Overall, the fold-change analysis indicated that the magnitude of qPCR-derived signal enhancement depended on both water matrix and wheat presence, with the largest increases observed for AC/PEG and EM in tap water with wheat.

### 3.2. ESV-normalized qPCR-derived viral signal

The ESV-normalized qPCR-derived viral signals, expressed as percentages of the corresponding non-enriched controls, are shown in Figure 2. This normalization provides a common relative scale for comparing sample-processing methods; however, the resulting percentages should not be interpreted as absolute viral recovery relative to the nominal spiked virus concentration.

**Figure 2.**
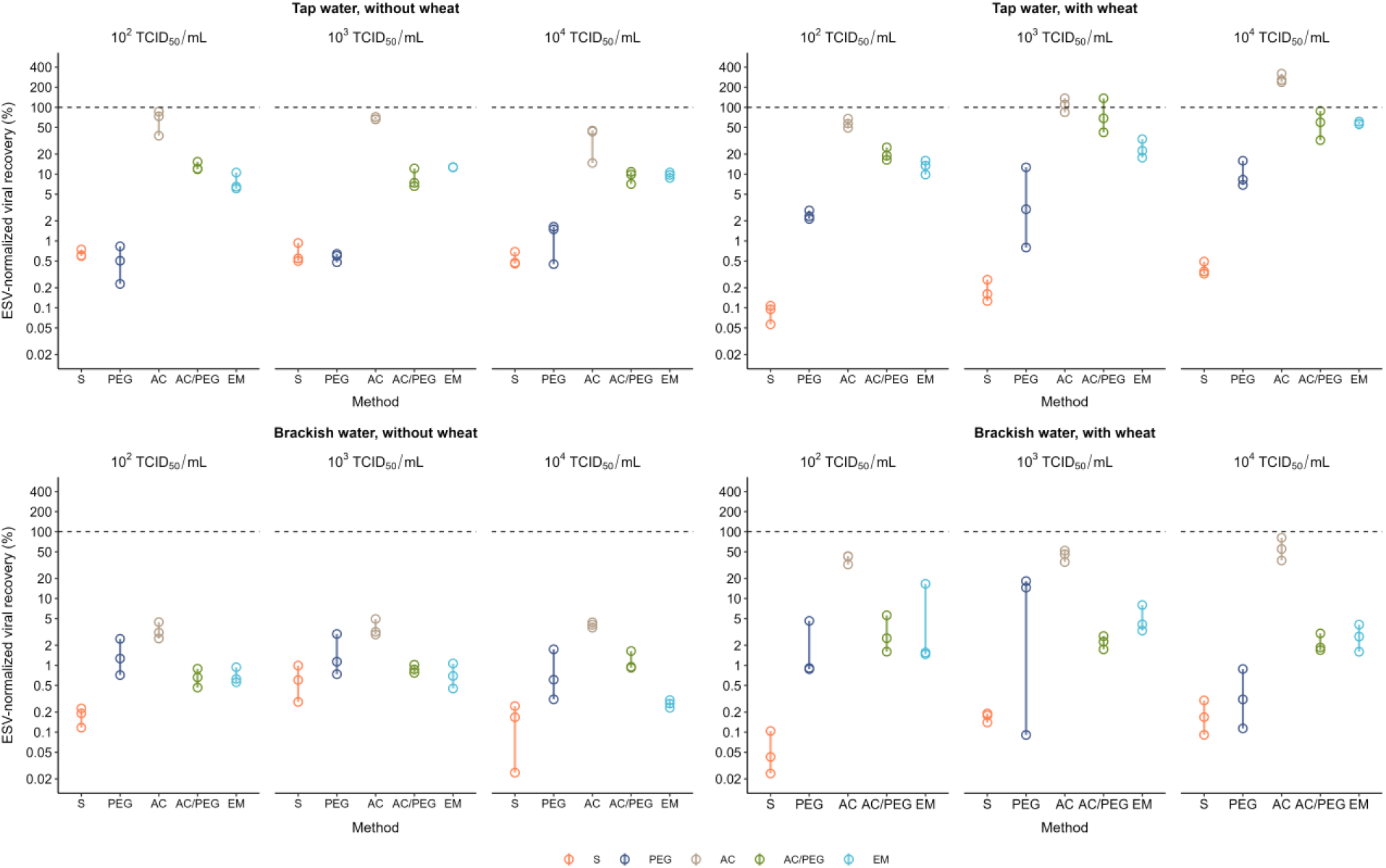
ESV-normalized qPCR-derived viral signal relative to the corresponding non-enriched control across sample-processing methods, water matrices, and wheat conditions. Individual points represent independent experimental replicates (n = 3 per concentration and method), and vertical lines represent the observed range across the three replicates. The dashed horizontal line indicates a signal equivalent to 100% of the ESV-normalized signal of the corresponding non-enriched control. The y-axis is shown on a base-10 logarithmic scale. Abbreviations: S, *swab sampling*; PEG, *polyethylene glycol precipitation*; AC, *activated carbon capture*; AC/PEG, *activated carbon capture* combined with PEG; EM, *electronegative membrane-based adsorption*.

AC consistently produced the highest mean ESV-normalized qPCR-derived viral signals across the different experimental conditions and spiked H4N6 concentrations. In tap water without wheat, AC reached mean ESV-normalized signals corresponding to 65.9, 68.2, and 34.4% of the respective non-enriched control values at 10², 10³, and 10⁴ TCID₅₀/mL, respectively. In the presence of wheat, the corresponding values were 58.0, 110.6, and 271.9%, respectively. In brackish water, AC likewise showed the highest mean ESV-normalized signals among the sample-processing methods, with values ranging from 3.4 to 4.0% without wheat and from 39.5 to 57.9% with wheat.

Among the other sample-processing methods, S generally showed the lowest ESV-normalized signals, particularly in tap water with wheat, whereas PEG, AC/PEG, and EM showed intermediate and more variable values depending on the water matrix and wheat condition. Overall, the ESV-normalized analysis consistently identified AC as the method yielding the highest relative qPCR-derived viral signal across the experimental conditions.

### 3.3. Effect of storage conditions on AIV RNA detection

AC/PEG and EM generally showed the strongest Ct-based performance across the experimental conditions. AC/PEG was therefore selected for Experiment 2 based on its overall performance and lower cost compared with EM. Preliminary screening of freshly collected brackish water samples provided additional support for its use (Supplementary Table S10).

The final model showed significant effects of water matrix, storage temperature, and storage duration (all *p* < 0.001), as well as significant interactions between water matrix and temperature, water matrix and storage duration, and temperature and storage duration (Supplementary Table S3).

Across both storage weeks and all storage temperatures, the qPCR-derived viral signal was consistently and significantly lower in brackish water than in tap water (Figure 3; Supplementary Table S6). Room-temperature storage resulted in a progressive decrease in the qPCR-derived viral signal from Week 2 to Week 4 in both matrices, with a greater loss observed in brackish water than in tap water (mean Ct increases of 2.7 and 1.3 cycles, respectively); the decrease was significant in both matrices (Supplementary Table S4). At 4 °C, the qPCR-derived viral signal decreased significantly in brackish water but not in tap water (Supplementary Table S4). At −20 °C, the qPCR-derived viral signal remained comparatively stable, with mean Ct increases of 0.8 and 1.0 cycles from Week 2 to Week 4 in tap and brackish water, respectively (Supplementary Table S4). Within each water matrix, the qPCR-derived viral signal was significantly lower at room temperature than at 4 °C or −20 °C at both Week 2 and Week 4. The qPCR-derived viral signal was also significantly lower at 4 °C than at −20 °C at both time points and in both matrices (Supplementary Table S5). The final model showed a high goodness of fit (R² = 0.985; adjusted R² = 0.979). Individual raw Ct values for all experimental conditions are provided in Supplementary Table S9.

**Figure 3.**
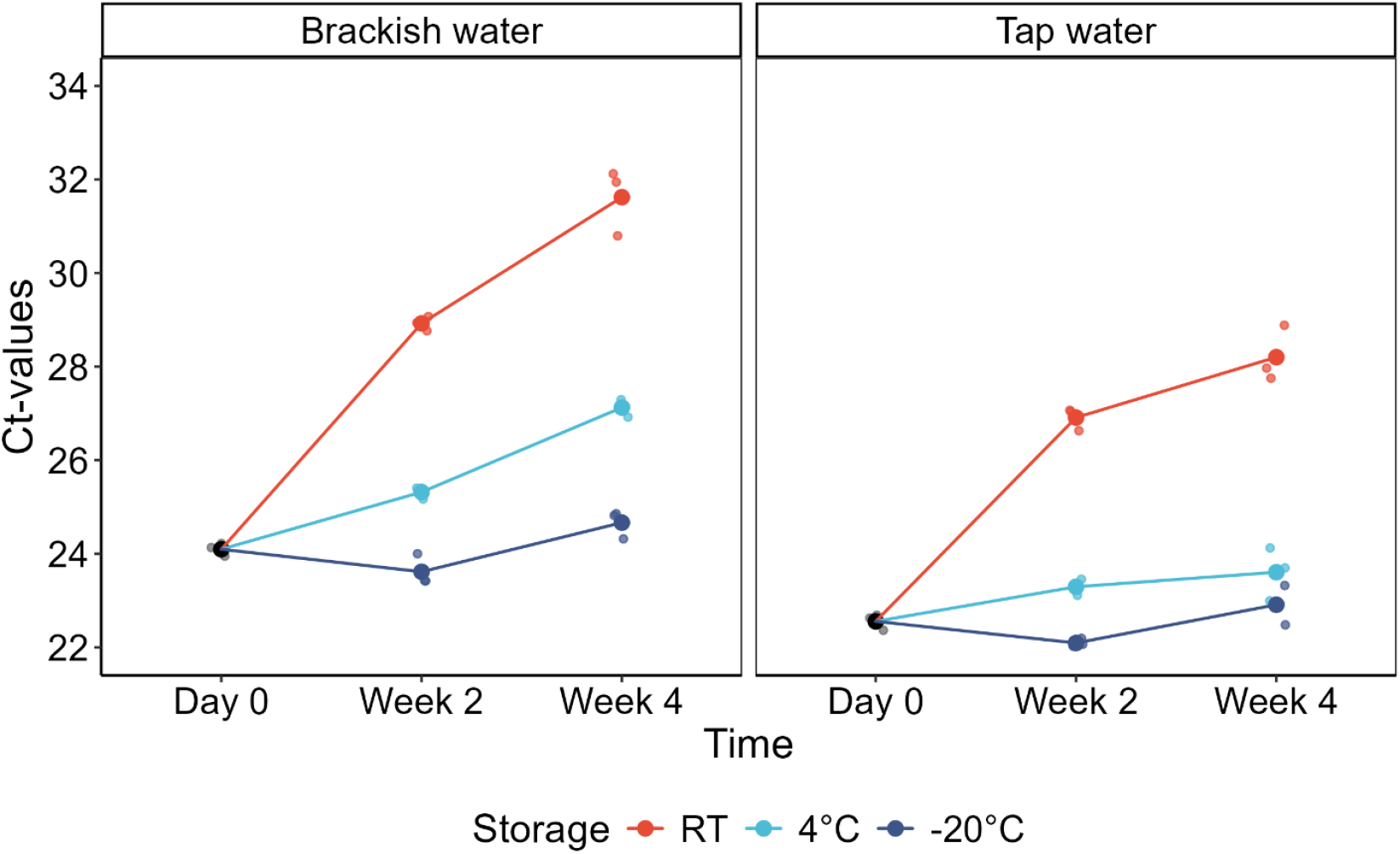
Effect of storage temperature on AIV RNA detection in 40-mL water samples spiked with H4N6 at 10³ TCID₅₀/mL and processed using AC/PEG over four weeks of storage. Points represent individual Ct values, with larger symbols indicating mean Ct values for each condition. Samples were stored at room temperature (RT), 4 °C, or −20 °C in tap water and brackish water and analyzed at Day 0, Week 2, and Week 4.

## Discussion

The present study evaluated five sample-processing methods for improving AIV RNA detection in low-volume environmental water samples under conditions designed to reflect the feed-baited water-bin surveillance approach, in which wheat-baited water was used to attract wild waterbirds (Rondi *et al*., 2026).

Abiotic factors, including temperature, pH, and salinity, can influence AIV persistence in water (Coombes *et al*., 2021). In the present study, the qPCR-derived viral signal was generally higher in tap water than in brackish water, potentially reflecting the difference in salinity between the two matrices (0.92 versus 7.3 ‰). Although the pH values were similar (7.1 and 6.9), initial water temperatures ranged from 21 to 24 °C, and a contribution of temperature to the observed differences cannot be excluded.

Beyond their potential effects on viral persistence, matrix characteristics may also influence the efficiency of virus enrichment and subsequent detection by RT-qPCR. Suspended and dissolved organic and inorganic material, for example, may affect virus adsorption and subsequent AIV RNA detection. Consistent with this, previous studies have shown that water physicochemical properties can affect viral recovery (Farkas *et al*., 2022) and that AIV recovery varies between surface-water sources (Deboosere *et al*., 2011). Similarly, Panzarin *et al*. (2025) evaluated the performance of different adsorbent materials for AIV recovery from both freshwater and brackish water, reporting differences in performance among materials and deployment times.

In the present study, the lower qPCR-derived viral signal observed with activated-carbon-based approaches in brackish water may reflect matrix-related effects on virus adsorption, potentially including competition from dissolved or suspended constituents for adsorption sites. Although this mechanism was not directly investigated, pre-filtration to reduce coarse particulate material before AC exposure could be explored as a potential optimization strategy.

The presence of wheat was also associated with a lower qPCR-derived viral signal, with the strongest reduction observed for swab sampling, particularly in tap water. This finding is especially relevant given that swabs were used for water sampling in Rondi *et al*. (2026). This reduction may reflect both physical and inhibitory effects of wheat-derived material. For example, wheat particles may physically accumulate on the swab material, potentially reducing its capacity to collect virus from the water. In addition, particulate or dissolved wheat-derived material carried into subsequent processing may also interfere with RNA extraction or RT-qPCR detection. Such interference may also explain the lower viral signal observed in the non-enriched control containing wheat than in the non-enriched control without wheat, in which samples were transferred directly to RNA extraction. This difference may have increased the apparent signal enhancement provided by the enrichment methods relative to the non-enriched control. In contrast, AC/PEG and EM showed comparatively high qPCR-derived viral signals, suggesting that these workflows were less affected by the presence of wheat. These explanations remain hypothetical, as the underlying mechanisms were not directly investigated in the present study.

Taken together, these findings have practical implications for feed-baited water-bin surveillance. Because wheat is an essential component of this approach and high viral loads are not necessarily expected in water collected from these bins, method performance in wheat-containing samples at low viral concentrations is particularly relevant. Under these conditions, AC/PEG yielded a higher fold-change in qPCR-derived viral signal than EM in tap water containing wheat at 10² TCID₅₀/mL (26.88 versus 18.68). Combined with its lower cost compared with EM, this suggests that AC/PEG may be a particularly suitable option for improving AIV RNA detection in low-volume samples collected using feed-baited water bins, particularly when tap water is used. Furthermore, preliminary screening of freshly collected brackish water samples, as described in *Materials and Methods*, provided additional evidence of the potential of AC/PEG for detecting AIV RNA in environmental water, with AIV RNA detected in 8 of 14 samples (57.1%; Supplementary Table S10). However, these samples were collected directly from the environment rather than from feed-baited water bins; therefore, the screening results do not directly validate the use of AC/PEG under the intended surveillance conditions.

AC showed the highest ESV-normalized qPCR-derived viral signals, whereas AC/PEG and EM generally showed the strongest performance based on Ct values. These differing rankings reflect the distinct information provided by the two metrics. ESV offers a complementary, volume-adjusted basis for comparing methods that differ in the amount of original sample represented in the RT-qPCR reaction, consistent with approaches used to compare virus enrichment methods in wastewater (Juel *et al*., 2021; Farkas *et al*., 2022). Therefore, the stronger performance of AC/PEG and EM based on Ct values and the higher ESV-normalized viral signal obtained with AC are not contradictory but reflect different aspects of workflow performance. The favourable ESV-normalized performance of AC, together with its ease of use and potential for scalable application reported previously (Holzer *et al*., 2025), highlights its potential as a simple and practical sample-processing approach. This is further supported by Gouthro *et al*. (2025), who detected influenza A and H5 RNA in surface waters using granular activated carbon-based passive samplers.

However, the relatively large eluate volume generated by AC represents a potential limitation for subsequent RT-qPCR analysis. Further optimization of the AC workflow could therefore focus on evaluating alternative approaches for concentrating the approximately 7-mL AC eluate, rather than relying exclusively on PEG, with the aim of increasing the viral signal available for RT-qPCR and improving analytical sensitivity.

For samples that cannot be processed immediately, the present results support storage at −20 °C for up to four weeks, with the qPCR-derived viral signal remaining relatively stable over the period investigated. This condition may be a practical alternative when −80 °C storage is not readily available. Nevertheless, samples should preferably be processed as soon as possible, and unnecessary temperature fluctuations and repeated freeze–thaw cycles should be avoided, as these may affect viral RNA integrity and detection (Ryba *et al*., 2012; Munguía-Ramírez *et al*., 2026).

Our results further indicate that storage at 4 °C, although resulting in significantly higher Ct values than storage at −20 °C, may represent a practical alternative where freezer facilities are unavailable, particularly for tap water samples. For brackish water, however, the more pronounced increase in Ct values indicates that refrigeration is less effective than freezing in maintaining the detectable viral signal. Previous studies have similarly demonstrated that storage temperature can influence viral RNA recovery from wastewater, although the direction and magnitude of these effects vary depending on sample characteristics and processing procedures (Hokajärvi *et al*., 2021; Islam *et al*., 2022). From an operational perspective, the availability of reliable refrigeration, freezer facilities and electricity represent an additional challenge for environmental pathogen surveillance in resource-limited settings (Truyens *et al*., 2025).

These findings should be interpreted in light of several limitations. The main comparative experiments were performed with laboratory-spiked water and a single H4N6 isolate, and therefore may not fully represent the variability encountered in naturally contaminated environmental samples or with other AIV subtypes. Only two water matrices were investigated, and potentially relevant matrix characteristics, such as organic and microbial content, were not systematically quantified. This limited our ability to identify the specific factors underlying the differences in qPCR-derived viral signal between water matrices. Furthermore, the recovery was assessed relative to the corresponding non-enriched control rather than as absolute recovery of the nominal amount of virus initially spiked. In addition, detection was based on viral RNA measured by RT-qPCR rather than on infectious virus. Accordingly, the results should be interpreted as relative differences in detectable AIV RNA rather than as absolute recovery of infectious AIV.

## Conclusions

Overall, AC/PEG and EM showed the strongest performance based on Ct values, although their relative performance varied across experimental conditions. Given its overall performance and lower cost compared with EM, AC/PEG may offer a practical option for feed-baited water-bin surveillance, particularly under the tap-water conditions evaluated. The qPCR-derived viral signal was generally higher in tap water than in brackish water, whereas wheat presence was associated with a lower signal, particularly for swab sampling. For storage, samples should preferably be processed as soon as possible, although storage at −20 °C maintained a relatively stable qPCR-derived viral signal for up to four weeks. Preliminary screening of freshly collected brackish water samples provided additional support for the use of AC/PEG, although further field evaluation under the intended surveillance conditions is needed.

## Data availability

The raw data underlying the results presented in this study are provided in Supplementary Tables S8–S10. No proprietary or restricted datasets were used.

## Acknowledgments

We would like to thank Simone Lüert, Anne Günther, Sharon Chepkwony, Arne Hegemann, Philipp Schwabl, Ann Charlott Keßler, and our colleagues at the Institute of International Animal Health/One Health (FLI) for their support and assistance.

## Supplementary material

**Supplementary Figure S1.**
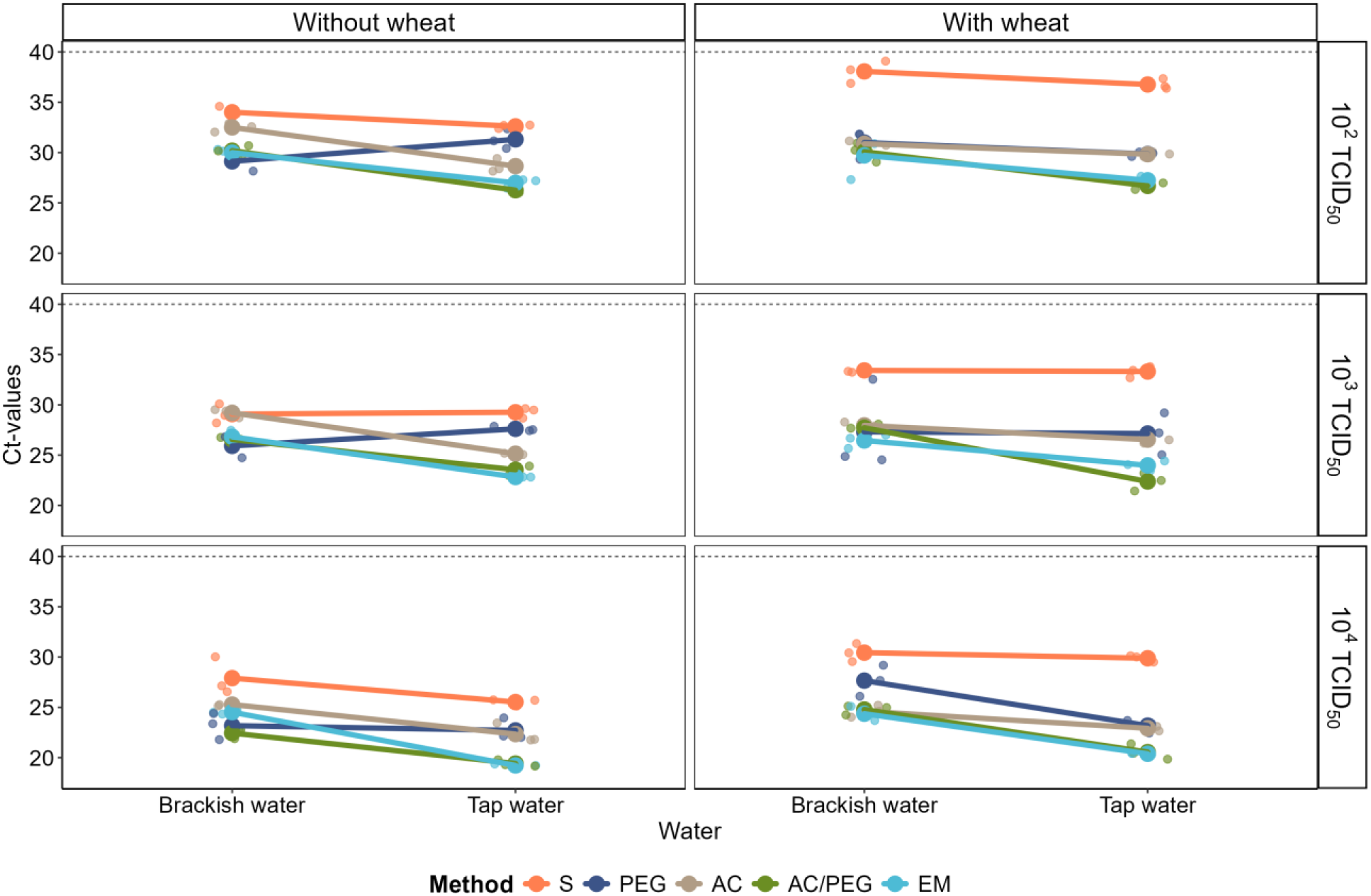
Ct values obtained for H4N6 after application of different sample-processing methods across water matrices (tap and brackish water), viral concentrations (10², 10^3^, 10⁴ TCID₅₀/mL), and wheat conditions (with or without wheat). The dashed horizontal line indicates the RT-qPCR cut-off value (Ct = 40). Abbreviations: S, *swab sampling*; PEG, *polyethylene glycol precipitation*; AC, *activated carbon capture*; AC/PEG, *activated carbon capture* combined with PEG; EM, *electronegative membrane-based adsorption*.

The five sample-processing methods showed variable Ct values across water matrices, spiked H4N6 concentrations, and wheat conditions (Supplementary Figure S1). The qPCR-derived viral signal generally increased with increasing spiked H4N6 concentration, although the magnitude and consistency of this relationship varied among methods and experimental conditions. Overall, the methods differed in their performance for AIV RNA detection.

For most methods, the qPCR-derived viral signal was lower in brackish than in tap water. S consistently yielded the lowest signal across water matrices, spiked H4N6 concentrations, and wheat conditions. Conversely, AC/PEG and EM generally yielded the highest signals, particularly in tap water. In brackish water, however, differences among methods were less consistent, and their relative ranking varied with spiked H4N6 concentration and wheat presence.

At 10² TCID₅₀/mL, differences among methods were readily apparent. In tap water, AC/PEG and EM consistently yielded the highest signals regardless of wheat presence, followed by AC, whereas PEG and particularly S yielded lower signals. In brackish water, differences were less pronounced. In the absence of wheat, PEG, AC/PEG, and EM showed similar signals, whereas no consistent differences among PEG, AC/PEG, EM, and AC were observed in the presence of wheat. S yielded the lowest signal under both wheat conditions.

At 10³ TCID₅₀/mL, the overall pattern in tap water remained similar, with AC/PEG and EM yielding the highest signals and S the lowest. In brackish water without wheat, signals were similar among PEG, EM, and AC/PEG, whereas AC consistently yielded lower values than these methods. In the presence of wheat, PEG showed greater variability than the other methods, while EM, AC/PEG, and AC exhibited comparatively narrower ranges of qPCR-derived viral signals. S again yielded the lowest signal.

At 10⁴ TCID₅₀/mL, differences among methods were generally less pronounced. In tap water, AC/PEG and EM again yielded the highest signals regardless of wheat presence. In brackish water without wheat, AC/PEG yielded the highest signal in two of the three replicates, whereas PEG yielded the highest signal in the remaining replicate. In the presence of wheat, the signals obtained with EM, AC/PEG, and AC were broadly similar, whereas PEG yielded lower values. S consistently yielded the lowest signal under all conditions.

**Supplementary Figure S2.**
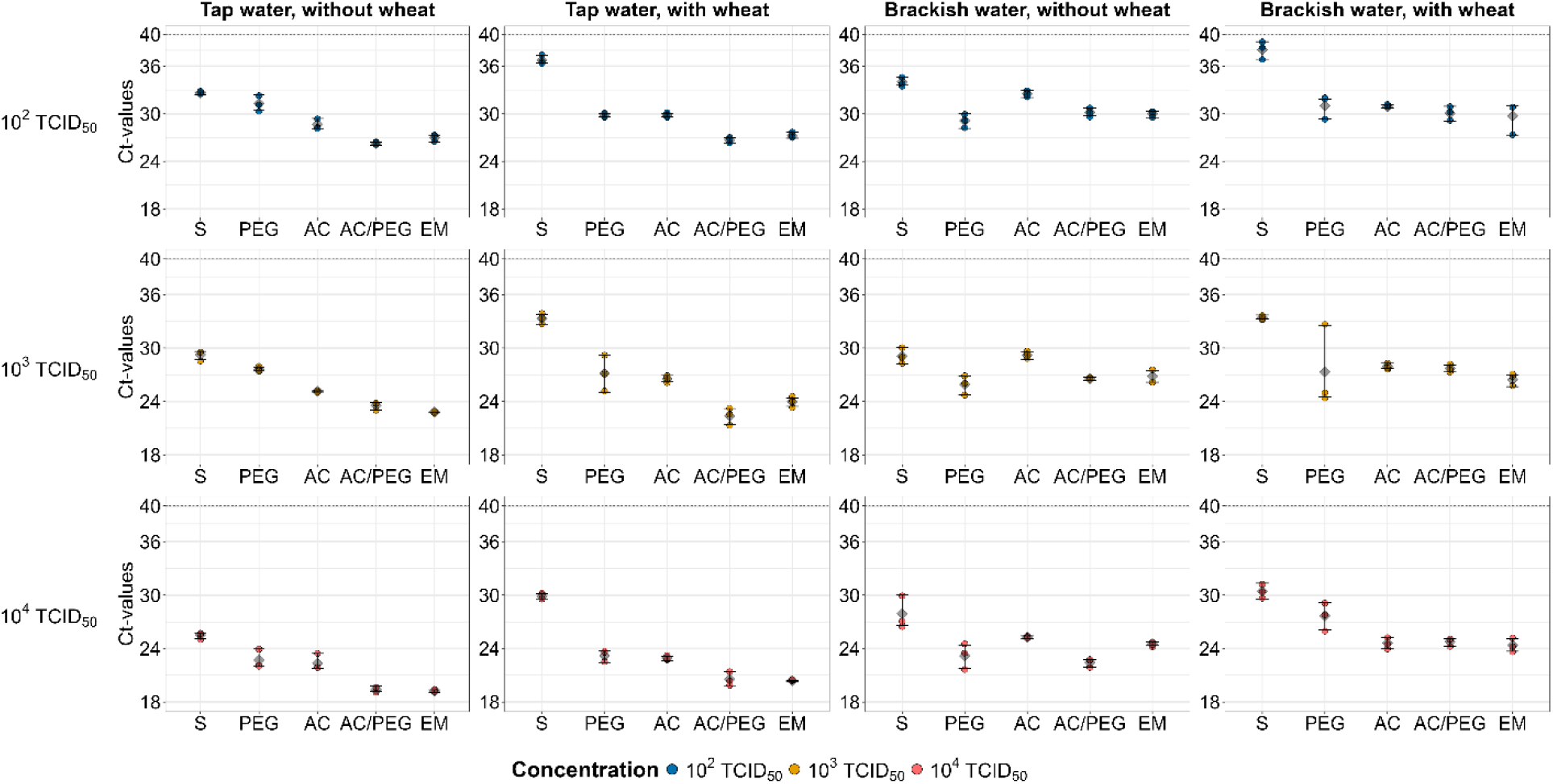
Individual replicate-level Ct values across sample-processing methods, water matrices, viral concentrations, and wheat conditions. Each panel represents one combination of water matrix and wheat condition, with rows corresponding to the three experimental viral concentrations (10², 10³, and 10⁴ TCID₅₀/mL). Individual points represent independent experimental replicates (n = 3 per sample-processing method and concentration). Large symbols indicate mean Ct values for each condition, with lines connecting the corresponding means across water matrices. The dashed horizontal line indicates the RT-qPCR cut-off value (Ct = 40). Abbreviations: S, *swab sampling*; PEG, *polyethylene glycol precipitation*; AC, *activated carbon capture*; AC/PEG, *activated carbon capture* combined with PEG; EM, *electronegative membrane-based adsorption*.

**Supplementary Table S1.** ANCOVA results for comparisons between sample-processing methods and non-enriched controls. β, model coefficient; CI, model-based 95% confidence interval; R², coefficient of determination; adjusted R², adjusted coefficient of determination. The interaction *p*-value refers to the concentration × method interaction in the full ANCOVA model. The interaction β and its confidence interval were obtained from the full ANCOVA model, regardless of whether the interaction term was retained in the final model. When this interaction was not significant (*p* ≥ 0.05), the reduced ANCOVA model without the interaction term was used as the final model; when the interaction was significant (*p* < 0.05), the full interaction model was retained. For models retaining the interaction, the Method β represents the difference in intercepts at log10(TCID₅₀/mL) = 0 and should not be interpreted as a concentration-specific effect. The interaction β represents the difference in regression slopes between the sample-processing method and the non-enriched control. Abbreviations: S, *swab sampling*; PEG, *polyethylene glycol precipitation*; AC, *activated carbon capture*; AC/PEG, *activated carbon capture* combined with PEG; EM, *electronegative membrane-based adsorption*.

| **Experimental condition** | **Sample-processing method** | **Final model** | **Interaction *p*** | **β<sub>2</sub> (Method)** | **95% CI** | ***p*** | **β<sub>3</sub> (Method × concentration)** | **95% CI** | **R²** | **Adjusted R²** |
|---|---|---|---|---|---|---|---|---|---|---|
| Tap water, without wheat | S | Reduced ANCOVA | 0.464 | 2.11 | 1.73 to 2.50 | <0.001 | 0.17 | -0.31 to 0.65 | 0.988 | 0.986 |
| Tap water, without wheat | PEG | Reduced ANCOVA | 0.151 | 0.19 | -0.52 to 0.90 | 0.575 | -0.59 | -1.43 to 0.24 | 0.963 | 0.958 |
| Tap water, without wheat | AC | Reduced ANCOVA | 0.102 | -1.64 | -2.23 to -1.05 | <0.001 | 0.56 | -0.13 to 1.24 | 0.968 | 0.963 |
| Tap water, without wheat | AC/PEG | Reduced ANCOVA | 0.302 | -3.94 | -4.41 to -3.47 | <0.001 | 0.29 | -0.29 to 0.86 | 0.985 | 0.984 |
| Tap water, without wheat | EM | Reduced ANCOVA | 0.427 | -4.01 | -4.37 to -3.64 | <0.001 | -0.17 | -0.62 to 0.28 | 0.992 | 0.991 |
| Tap water, with wheat | S | Interaction model | <0.001 | 7.37 | 5.59 to 9.14 | <0.001 | -1.14 | -1.71 to -0.57 | 0.983 | 0.979 |
| Tap water, with wheat | PEG | Reduced ANCOVA | 0.073 | -2.65 | -3.66 to -1.64 | <0.001 | -1.04 | -2.18 to 0.11 | 0.893 | 0.878 |
| Tap water, with wheat | AC | Interaction model | <0.001 | 0.53 | -0.97 to 2.03 | 0.461 | -1.16 | -1.64 to -0.68 | 0.985 | 0.982 |
| Tap water, with wheat | AC/PEG | Reduced ANCOVA | 0.094 | -6.17 | -6.97 to -5.38 | <0.001 | -0.77 | -1.68 to 0.15 | 0.965 | 0.960 |
| Tap water, with wheat | EM | Interaction model | <0.001 | -2.13 | -3.72 to -0.54 | 0.012 | -1.13 | -1.64 to -0.62 | 0.990 | 0.988 |
| Brackish water, without wheat | S | Reduced ANCOVA | 0.524 | 3.71 | 2.64 to 4.79 | <0.001 | 0.41 | -0.94 to 1.76 | 0.917 | 0.906 |
| Brackish water, without wheat | PEG | Reduced ANCOVA | 0.279 | -0.54 | -1.31 to 0.22 | 0.151 | 0.49 | -0.44 to 1.43 | 0.935 | 0.926 |
| Brackish water, without wheat | AC | Reduced ANCOVA | 0.421 | 2.37 | 2.06 to 2.69 | <0.001 | -0.15 | -0.54 to 0.24 | 0.992 | 0.991 |
| Brackish water, without wheat | AC/PEG | Reduced ANCOVA | 0.053 | -0.22 | -0.60 to 0.15 | 0.221 | -0.41 | -0.82 to 0.01 | 0.987 | 0.986 |
| Brackish water, without wheat | EM | Interaction model | 0.005 | -1.69 | -3.16 to -0.22 | 0.027 | 0.73 | 0.26 to 1.20 | 0.983 | 0.979 |
| Brackish water, with wheat | S | Reduced ANCOVA | 0.076 | 4.61 | 3.74 to 5.47 | <0.001 | -0.88 | -1.86 to 0.10 | 0.954 | 0.948 |
| Brackish water, with wheat | PEG | Reduced ANCOVA | 0.312 | -0.72 | -2.82 to 1.39 | 0.480 | 1.27 | -1.32 to 3.85 | 0.500 | 0.433 |
| Brackish water, with wheat | AC | Reduced ANCOVA | 0.557 | -1.55 | -2.10 to -1.01 | <0.001 | -0.19 | -0.88 to 0.50 | 0.965 | 0.960 |
| Tap water, without wheat | S | Reduced ANCOVA | 0.464 | 2.11 | 1.73 to 2.50 | <0.001 | 0.17 | -0.31 to 0.65 | 0.988 | 0.986 |
| Tap water, without wheat | PEG | Reduced ANCOVA | 0.151 | 0.19 | -0.52 to 0.90 | 0.575 | -0.59 | -1.43 to 0.24 | 0.963 | 0.958 |
| Tap water, without wheat | AC | Reduced ANCOVA | 0.102 | -1.64 | -2.23 to -1.05 | <0.001 | 0.56 | -0.13 to 1.24 | 0.968 | 0.963 |
| Brackish water, with wheat | AC/PEG | Reduced ANCOVA | 0.455 | -1.84 | -2.49 to -1.20 | <0.001 | 0.29 | -0.52 to 1.09 | 0.946 | 0.939 |
| Brackish water, with wheat | EM | Reduced ANCOVA | 0.668 | -2.52 | -3.52 to -1.52 | <0.001 | 0.26 | -1.01 to 1.52 | 0.893 | 0.878 |

**Supplementary Table S2.** Concentration-specific planned comparisons for sample-processing methods showing a significant concentration × method interaction. SE, standard error; df, degrees of freedom; CI, model-based 95% confidence interval. Estimated Ct differences were obtained from estimated marginal means of the final ANCOVA model and represent the model-estimated difference between the indicated sample-processing method and the corresponding non-enriched control at the specified viral concentration. Negative values indicate lower Ct values for the sample-processing method than for the non-enriched control, whereas positive values indicate higher Ct values. Comparisons were evaluated at the three pre-specified experimental concentrations (10², 10³, and 10⁴ TCID₅₀/mL). These concentration-specific comparisons were defined *a priori* as planned comparisons to characterize the interaction across the experimental concentration range. Abbreviations: S, *swab sampling*; AC, *activated carbon capture*; EM, *electronegative membrane-based adsorption*.

| **Experimental condition** | **Sample-processing method** | **Viral concentration (TCID<sub>50</sub>/mL)** | **Estimated Ct difference** | **SE** | **df** | **95% CI** | ***p* (Holm-adjusted)** |
|---|---|---|---|---|---|---|---|
| Tap water, with wheat | S | 10<sup>2</sup> | 5.08 | 0.34 | 14.0 | 4.34 to 5.82 | <0.001 |
| Tap water, with wheat | S | 10<sup>3</sup> | 3.94 | 0.22 | 14.0 | 3.47 to 4.40 | <0.001 |
| Tap water, with wheat | S | 10<sup>4</sup> | 2.79 | 0.34 | 14.0 | 2.06 to 3.53 | <0.001 |
| Tap water, with wheat | AC | 10<sup>2</sup> | -1.79 | 0.29 | 14.0 | -2.41 to -1.17 | <0.001 |
| Tap water, with wheat | AC | 10<sup>3</sup> | -2.95 | 0.18 | 14.0 | -3.34 to -2.56 | <0.001 |
| Tap water, with wheat | AC | 10<sup>4</sup> | -4.11 | 0.29 | 14.0 | -4.73 to -3.49 | <0.001 |
| Tap water, with wheat | EM | 10<sup>2</sup> | -4.38 | 0.31 | 14.0 | -5.04 to -3.72 | <0.001 |
| Tap water, with wheat | EM | 10<sup>3</sup> | -5.51 | 0.19 | 14.0 | -5.93 to -5.09 | <0.001 |
| Tap water, with wheat | EM | 10<sup>4</sup> | -6.64 | 0.31 | 14.0 | -7.30 to -5.98 | <0.001 |
| Brackish water, without wheat | EM | 10<sup>2</sup> | -0.24 | 0.28 | 14.0 | -0.85 to 0.37 | 0.421 |
| Brackish water, without wheat | EM | 10<sup>3</sup> | 0.49 | 0.18 | 14.0 | 0.11 to 0.88 | 0.032 |
| Brackish water, without wheat | EM | 10<sup>4</sup> | 1.22 | 0.28 | 14.0 | 0.61 to 1.83 | 0.002 |

**Supplementary Table S3.** Type III ANOVA results for the final linear model of Ct values. df, degrees of freedom. The final model included Water, Temperature, Week, and their two-way interactions. Day 0 samples were excluded from the factorial analysis because no storage temperature was applicable.

| **Effect** | **df** | **F** | ***p*** |
|---|---|---|---|
| Water | 1 | 310.20 | <0.001 |
| Temperature | 2 | 613.43 | <0.001 |
| Week | 1 | 97.08 | <0.001 |
| Water × Temperature | 2 | 7.48 | 0.003 |
| Water × Week | 1 | 15.00 | <0.001 |
| Temperature × Week | 2 | 6.16 | 0.006 |
| Residuals | 26 | — | — |

**Supplementary Table S4.**
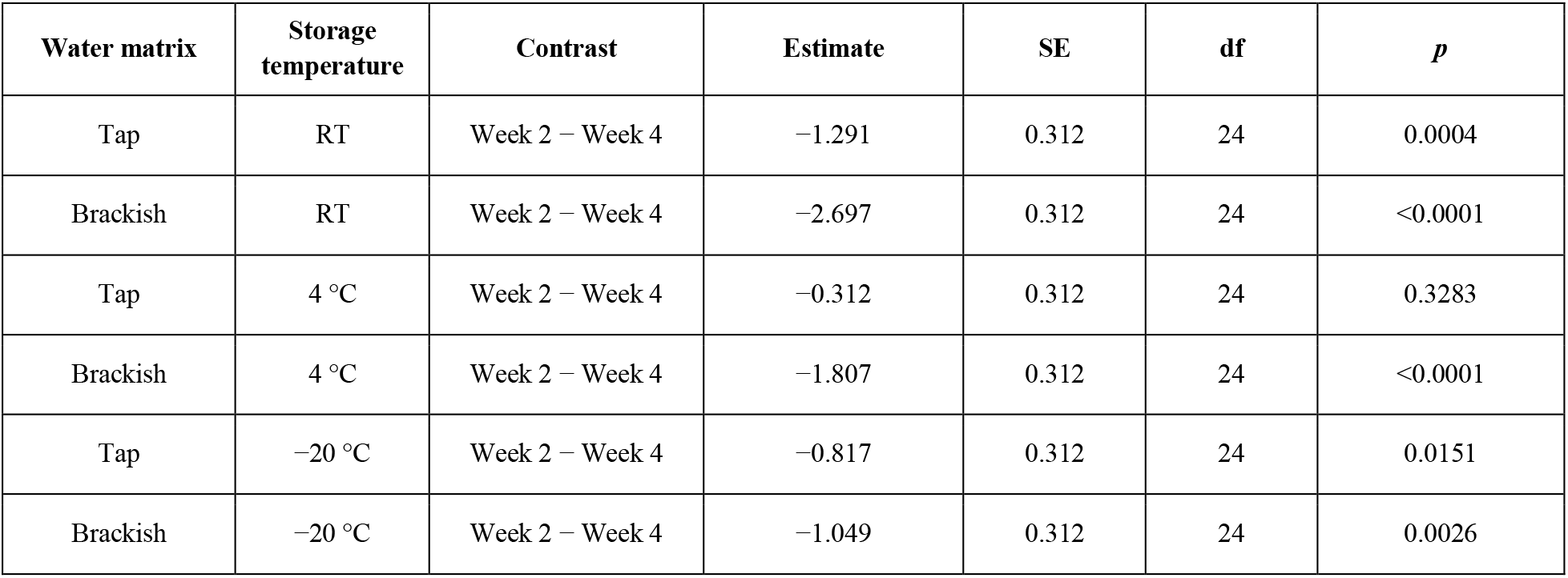
Pairwise comparisons of Ct values between Week 2 and Week 4 within each water matrix and storage temperature. SE, standard error; df, degrees of freedom. Estimated Ct differences were obtained from estimated marginal means of the final factorial linear model and represent the model-estimated difference between Week 2 and Week 4 within the indicated water matrix and storage temperature. Negative values indicate lower Ct values at Week 2 than at Week 4, whereas positive values indicate higher Ct values at Week 2 than at Week 4. Comparisons were performed separately for each combination of water matrix and storage temperature. *P*-values were adjusted using Tukey’s method. Abbreviations: RT, room temperature.

| **Water matrix** | **Storage temperature** | **Contrast** | **Estimate** | **SE** | **df** | ***p*** |
|---|---|---|---|---|---|---|
| Tap | RT | Week 2 − Week 4 | −1.291 | 0.312 | 24 | 0.0004 |
| Brackish | RT | Week 2 − Week 4 | −2.697 | 0.312 | 24 | <0.0001 |
| Tap | 4 °C | Week 2 − Week 4 | −0.312 | 0.312 | 24 | 0.3283 |
| Brackish | 4 °C | Week 2 − Week 4 | −1.807 | 0.312 | 24 | <0.0001 |
| Tap | −20 °C | Week 2 − Week 4 | −0.817 | 0.312 | 24 | 0.0151 |
| Brackish | −20 °C | Week 2 − Week 4 | −1.049 | 0.312 | 24 | 0.0026 |

**Supplementary Table S5.**
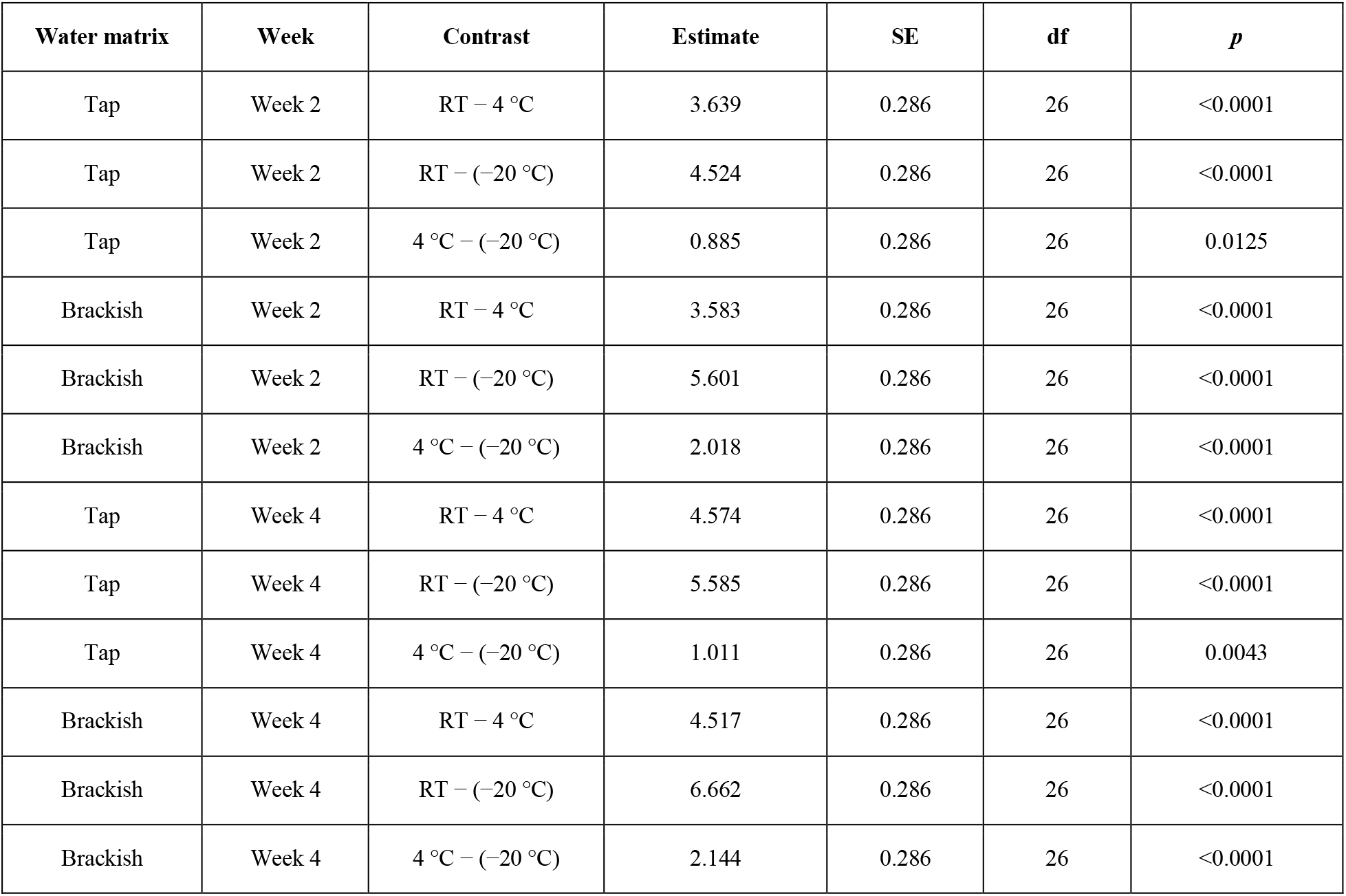
Pairwise comparisons of Ct values among storage temperatures within each water matrix and storage week. SE, standard error; df, degrees of freedom. Estimated Ct differences were obtained from estimated marginal means of the final factorial linear model and represent the model-estimated difference between the indicated storage temperatures within the specified water matrix and storage week. Positive values indicate higher Ct values for the first temperature in the contrast, whereas negative values indicate lower Ct values for the first temperature. Comparisons were performed separately for each combination of water matrix and storage week. *P*-values were adjusted using Tukey’s method. Abbreviations: RT, room temperature.

| **Water matrix** | **Week** | **Contrast** | **Estimate** | **SE** | **df** | ***p*** |
|---|---|---|---|---|---|---|
| Tap | Week 2 | RT − 4 °C | 3.639 | 0.286 | 26 | <0.0001 |
| Tap | Week 2 | RT − (−20 °C) | 4.524 | 0.286 | 26 | <0.0001 |
| Tap | Week 2 | 4 °C − (−20 °C) | 0.885 | 0.286 | 26 | 0.0125 |
| Brackish | Week 2 | RT − 4 °C | 3.583 | 0.286 | 26 | <0.0001 |
| Brackish | Week 2 | RT − (−20 °C) | 5.601 | 0.286 | 26 | <0.0001 |
| Brackish | Week 2 | 4 °C − (−20 °C) | 2.018 | 0.286 | 26 | <0.0001 |
| Tap | Week 4 | RT − 4 °C | 4.574 | 0.286 | 26 | <0.0001 |
| Tap | Week 4 | RT − (−20 °C) | 5.585 | 0.286 | 26 | <0.0001 |
| Tap | Week 4 | 4 °C − (−20 °C) | 1.011 | 0.286 | 26 | 0.0043 |
| Brackish | Week 4 | RT − 4 °C | 4.517 | 0.286 | 26 | <0.0001 |
| Brackish | Week 4 | RT − (−20 °C) | 6.662 | 0.286 | 26 | <0.0001 |
| Brackish | Week 4 | 4 °C − (−20 °C) | 2.144 | 0.286 | 26 | <0.0001 |

**Supplementary Table S6.** Pairwise comparisons of Ct values between tap and brackish water within each storage temperature and storage week. SE, standard error; df, degrees of freedom. Estimated Ct differences were obtained from estimated marginal means of the final factorial linear model and represent the model-estimated difference between tap and brackish water within the specified storage temperature and storage week. Negative values indicate lower Ct values in tap water than in brackish water, whereas positive values indicate higher Ct values in tap water than in brackish water. Comparisons were performed separately for each combination of storage temperature and storage week. *P*-values were adjusted using Tukey’s method. Abbreviations: RT, room temperature.

| **Storage temperature** | **Week** | **Contrast** | **Estimate** | **SE** | **df** | **t-ratio** | ***p*** |
|---|---|---|---|---|---|---|---|
| RT | Week 2 | Tap − Bodden | −2.19 | 0.27 | 26 | −8.131 | <0.0001 |
| 4 °C | Week 2 | Tap − Bodden | −2.25 | 0.27 | 26 | −8.340 | <0.0001 |
| −20 °C | Week 2 | Tap − Bodden | −1.12 | 0.27 | 26 | −4.138 | 0.0003 |
| RT | Week 4 | Tap − Bodden | −3.24 | 0.27 | 26 | −12.004 | <0.0001 |
| 4 °C | Week 4 | Tap − Bodden | −3.29 | 0.27 | 26 | −12.213 | <0.0001 |
| −20 °C | Week 4 | Tap − Bodden | −2.16 | 0.27 | 26 | −8.011 | <0.0001 |

**Supplementary Table S7.** Ct values obtained from the H4N6 standard curve at each nominal TCID₅₀ concentration. Abbreviation: NA, not detected. The RT-qPCR cut-off value was set at Ct = 40.

| **Nominal concentration (TCID<sub>50</sub>/mL)** | **Replicate 1 Ct** | **Replicate 2 Ct** | **Replicate 3 Ct** | **Replicate 4 Ct** | **Mean Ct** |
|---|---|---|---|---|---|
| 10<sup>−1</sup> | NA | NA | NA | NA | - |
| 10<sup>0</sup> | 38.77 | 39.78 | 39.79 | 39.12 | 39.37 |
| 10<sup>1</sup> | 34.16 | 34.20 | 34.14 | 34.95 | 34.36 |
| 10<sup>2</sup> | 31.40 | 31.33 | 31.44 | 31.10 | 31.32 |
| 10<sup>3</sup> | 28.69 | 27.93 | 27.68 | 28.17 | 28.12 |
| 10<sup>4</sup> | 24.24 | 24.31 | 24.43 | 24.11 | 24.27 |
| 10<sup>5</sup> | 20.54 | 20.64 | 20.73 | 20.71 | 20.66 |
| 10<sup>6</sup> | 19.09 | 18.58 | 17.71 | 18.32 | 18.43 |

**Supplementary Table S8.** Individual Ct values obtained from H4N6-spiked water samples processed using the five sample-processing methods across two water matrices (tap and brackish), three viral concentrations (10², 10³ and 10⁴ TCID₅₀/mL), and the presence or absence of wheat. Abbreviations: S, *swab sampling*; PEG, *polyethylene glycol precipitation*; AC, *activated carbon capture*; AC/PEG, *activated carbon capture* combined with PEG; EM, *electronegative membrane-based adsorption*.

| **Experimental condition** | **Sample-processing method** | **Viral concentration (TCID<sub>50</sub>/mL)** | **Replicate** | **Ct value** |
|---|---|---|---|---|
| Tap water, without wheat | S | 10<sup>2</sup> | 1 | 32.70 |
| Tap water, without wheat | S | 10<sup>2</sup> | 2 | 32.73 |
| Tap water, without wheat | S | 10<sup>2</sup> | 3 | 32.39 |
| Tap water, without wheat | PEG | 10<sup>2</sup> | 1 | 32.36 |
| Tap water, without wheat | PEG | 10<sup>2</sup> | 2 | 31.16 |
| Tap water, without wheat | PEG | 10<sup>2</sup> | 3 | 30.41 |
| Tap water, without wheat | AC | 10<sup>2</sup> | 1 | 28.40 |
| Tap water, without wheat | AC | 10<sup>2</sup> | 2 | 29.41 |
| Tap water, without wheat | AC | 10<sup>2</sup> | 3 | 28.15 |
| Tap water, without wheat | AC/PEG | 10<sup>2</sup> | 1 | 26.40 |
| Tap water, without wheat | AC/PEG | 10<sup>2</sup> | 2 | 26.01 |
| Tap water, without wheat | AC/PEG | 10<sup>2</sup> | 3 | 26.37 |
| Tap water, without wheat | EM | 10<sup>2</sup> | 1 | 27.20 |
| Tap water, without wheat | EM | 10<sup>2</sup> | 2 | 27.30 |
| Tap water, without wheat | EM | 10<sup>2</sup> | 3 | 26.47 |
| Tap water, without wheat | S | 10<sup>3</sup> | 1 | 29.48 |
| Tap water, without wheat | S | 10<sup>3</sup> | 2 | 29.62 |
| Tap water, without wheat | S | 10<sup>3</sup> | 3 | 28.68 |
| Tap water, without wheat | PEG | 10<sup>3</sup> | 1 | 27.53 |
| Tap water, without wheat | PEG | 10<sup>3</sup> | 2 | 27.87 |
| Tap water, without wheat | PEG | 10<sup>3</sup> | 3 | 27.43 |
| Tap water, without wheat | AC | 10<sup>3</sup> | 1 | 25.18 |
| Tap water, without wheat | AC | 10<sup>3</sup> | 2 | 25.06 |
| Tap water, without wheat | AC | 10<sup>3</sup> | 3 | 25.19 |
| Tap water, without wheat | AC/PEG | 10<sup>3</sup> | 1 | 23.74 |
| Tap water, without wheat | AC/PEG | 10<sup>3</sup> | 2 | 23.91 |
| Tap water, without wheat | AC/PEG | 10<sup>3</sup> | 3 | 22.99 |
| Tap water, without wheat | EM | 10<sup>3</sup> | 1 | 22.81 |
| Tap water, without wheat | EM | 10<sup>3</sup> | 2 | 22.83 |
| Tap water, without wheat | EM | 10<sup>3</sup> | 3 | 22.83 |
| Tap water, without wheat | S | 10<sup>4</sup> | 1 | 25.70 |
| Tap water, without wheat | S | 10<sup>4</sup> | 2 | 25.13 |
| Tap water, without wheat | S | 10<sup>4</sup> | 3 | 25.75 |
| Tap water, without wheat | PEG | 10<sup>4</sup> | 1 | 22.01 |
| Tap water, without wheat | PEG | 10<sup>4</sup> | 2 | 22.15 |
| Tap water, without wheat | PEG | 10<sup>4</sup> | 3 | 23.96 |
| Tap water, without wheat | AC | 10<sup>4</sup> | 1 | 21.82 |
| Tap water, without wheat | AC | 10<sup>4</sup> | 2 | 21.76 |
| Tap water, without wheat | AC | 10<sup>4</sup> | 3 | 23.45 |
| Tap water, without wheat | AC/PEG | 10<sup>4</sup> | 1 | 19.16 |
| Tap water, without wheat | AC/PEG | 10<sup>4</sup> | 2 | 19.30 |
| Tap water, without wheat | AC/PEG | 10<sup>4</sup> | 3 | 19.79 |
| Tap water, without wheat | EM | 10<sup>4</sup> | 1 | 19.22 |
| Tap water, without wheat | EM | 10<sup>4</sup> | 2 | 19.37 |
| Tap water, without wheat | EM | 10<sup>4</sup> | 3 | 19.09 |
| Tap water, with wheat | S | 10<sup>2</sup> | 1 | 37.35 |
| Tap water, with wheat | S | 10<sup>2</sup> | 2 | 36.57 |
| Tap water, with wheat | S | 10<sup>2</sup> | 3 | 36.38 |
| Tap water, with wheat | PEG | 10<sup>2</sup> | 1 | 29.92 |
| Tap water, with wheat | PEG | 10<sup>2</sup> | 2 | 30.05 |
| Tap water, with wheat | PEG | 10<sup>2</sup> | 3 | 29.61 |
| Tap water, with wheat | AC | 10<sup>2</sup> | 1 | 30.07 |
| Tap water, with wheat | AC | 10<sup>2</sup> | 2 | 29.59 |
| Tap water, with wheat | AC | 10<sup>2</sup> | 3 | 29.85 |
| Tap water, with wheat | AC/PEG | 10<sup>2</sup> | 1 | 26.33 |
| Tap water, with wheat | AC/PEG | 10<sup>2</sup> | 2 | 26.98 |
| Tap water, with wheat | AC/PEG | 10<sup>2</sup> | 3 | 26.76 |
| Tap water, with wheat | EM | 10<sup>2</sup> | 1 | 27.64 |
| Tap water, with wheat | EM | 10<sup>2</sup> | 2 | 27.17 |
| Tap water, with wheat | EM | 10<sup>2</sup> | 3 | 26.92 |
| Tap water, with wheat | S | 10<sup>3</sup> | 1 | 33.43 |
| Tap water, with wheat | S | 10<sup>3</sup> | 2 | 32.69 |
| Tap water, with wheat | S | 10<sup>3</sup> | 3 | 33.80 |
| Tap water, with wheat | PEG | 10<sup>3</sup> | 1 | 25.03 |
| Tap water, with wheat | PEG | 10<sup>3</sup> | 2 | 29.20 |
| Tap water, with wheat | PEG | 10<sup>3</sup> | 3 | 27.21 |
| Tap water, with wheat | AC | 10<sup>3</sup> | 1 | 26.51 |
| Tap water, with wheat | AC | 10<sup>3</sup> | 2 | 26.92 |
| Tap water, with wheat | AC | 10<sup>3</sup> | 3 | 26.19 |
| Tap water, with wheat | AC/PEG | 10<sup>3</sup> | 1 | 22.48 |
| Tap water, with wheat | AC/PEG | 10<sup>3</sup> | 2 | 23.21 |
| Tap water, with wheat | AC/PEG | 10<sup>3</sup> | 3 | 21.44 |
| Tap water, with wheat | EM | 10<sup>3</sup> | 1 | 23.46 |
| Tap water, with wheat | EM | 10<sup>3</sup> | 2 | 24.06 |
| Tap water, with wheat | EM | 10<sup>3</sup> | 3 | 24.42 |
| Tap water, with wheat | S | 10<sup>4</sup> | 1 | 30.13 |
| Tap water, with wheat | S | 10<sup>4</sup> | 2 | 30.01 |
| Tap water, with wheat | S | 10<sup>4</sup> | 3 | 29.50 |
| Tap water, with wheat | PEG | 10<sup>4</sup> | 1 | 23.70 |
| Tap water, with wheat | PEG | 10<sup>4</sup> | 2 | 22.44 |
| Tap water, with wheat | PEG | 10<sup>4</sup> | 3 | 23.42 |
| Tap water, with wheat | AC | 10<sup>4</sup> | 1 | 23.10 |
| Tap water, with wheat | AC | 10<sup>4</sup> | 2 | 22.66 |
| Tap water, with wheat | AC | 10<sup>4</sup> | 3 | 22.99 |
| Tap water, with wheat | AC/PEG | 10<sup>4</sup> | 1 | 21.37 |
| Tap water, with wheat | AC/PEG | 10<sup>4</sup> | 2 | 20.44 |
| Tap water, with wheat | AC/PEG | 10<sup>4</sup> | 3 | 19.85 |
| Tap water, with wheat | EM | 10<sup>4</sup> | 1 | 20.30 |
| Tap water, with wheat | EM | 10<sup>4</sup> | 2 | 20.44 |
| Tap water, with wheat | EM | 10<sup>4</sup> | 3 | 20.42 |
| Brackish water, without wheat | S | 10<sup>2</sup> | 1 | 33.59 |
| Brackish water, without wheat | S | 10<sup>2</sup> | 2 | 33.85 |
| Brackish water, without wheat | S | 10<sup>2</sup> | 3 | 34.59 |
| Brackish water, without wheat | PEG | 10<sup>2</sup> | 1 | 28.16 |
| Brackish water, without wheat | PEG | 10<sup>2</sup> | 2 | 30.04 |
| Brackish water, without wheat | PEG | 10<sup>2</sup> | 3 | 29.18 |
| Brackish water, without wheat | AC | 10<sup>2</sup> | 1 | 32.89 |
| Brackish water, without wheat | AC | 10<sup>2</sup> | 2 | 32.58 |
| Brackish water, without wheat | AC | 10<sup>2</sup> | 3 | 32.04 |
| Brackish water, without wheat | AC/PEG | 10<sup>2</sup> | 1 | 30.69 |
| Brackish water, without wheat | AC/PEG | 10<sup>2</sup> | 2 | 29.71 |
| Brackish water, without wheat | AC/PEG | 10<sup>2</sup> | 3 | 30.16 |
| Brackish water, without wheat | EM | 10<sup>2</sup> | 1 | 30.14 |
| Brackish water, without wheat | EM | 10<sup>2</sup> | 2 | 29.53 |
| Brackish water, without wheat | EM | 10<sup>2</sup> | 3 | 30.31 |
| Brackish water, without wheat | S | 10<sup>3</sup> | 1 | 28.95 |
| Brackish water, without wheat | S | 10<sup>3</sup> | 2 | 30.09 |
| Brackish water, without wheat | S | 10<sup>3</sup> | 3 | 28.20 |
| Brackish water, without wheat | PEG | 10<sup>3</sup> | 1 | 26.18 |
| Brackish water, without wheat | PEG | 10<sup>3</sup> | 2 | 24.74 |
| Brackish water, without wheat | PEG | 10<sup>3</sup> | 3 | 26.83 |
| Brackish water, without wheat | AC | 10<sup>3</sup> | 1 | 29.52 |
| Brackish water, without wheat | AC | 10<sup>3</sup> | 2 | 29.38 |
| Brackish water, without wheat | AC | 10<sup>3</sup> | 3 | 28.71 |
| Brackish water, without wheat | AC/PEG | 10<sup>3</sup> | 1 | 26.76 |
| Brackish water, without wheat | AC/PEG | 10<sup>3</sup> | 2 | 26.58 |
| Brackish water, without wheat | AC/PEG | 10<sup>3</sup> | 3 | 26.34 |
| Brackish water, without wheat | EM | 10<sup>3</sup> | 1 | 26.17 |
| Brackish water, without wheat | EM | 10<sup>3</sup> | 2 | 26.82 |
| Brackish water, without wheat | EM | 10<sup>3</sup> | 3 | 27.47 |
| Brackish water, without wheat | S | 10<sup>4</sup> | 1 | 30.02 |
| Brackish water, without wheat | S | 10<sup>4</sup> | 2 | 27.14 |
| Brackish water, without wheat | S | 10<sup>4</sup> | 3 | 26.56 |
| Brackish water, without wheat | PEG | 10<sup>4</sup> | 1 | 23.37 |
| Brackish water, without wheat | PEG | 10<sup>4</sup> | 2 | 24.39 |
| Brackish water, without wheat | PEG | 10<sup>4</sup> | 3 | 21.79 |
| Brackish water, without wheat | AC | 10<sup>4</sup> | 1 | 25.42 |
| Brackish water, without wheat | AC | 10<sup>4</sup> | 2 | 25.27 |
| Brackish water, without wheat | AC | 10<sup>4</sup> | 3 | 25.14 |
| Brackish water, without wheat | AC/PEG | 10<sup>4</sup> | 1 | 22.75 |
| Brackish water, without wheat | AC/PEG | 10<sup>4</sup> | 2 | 21.88 |
| Brackish water, without wheat | AC/PEG | 10<sup>4</sup> | 3 | 22.70 |
| Brackish water, without wheat | EM | 10<sup>4</sup> | 1 | 24.33 |
| Brackish water, without wheat | EM | 10<sup>4</sup> | 2 | 24.51 |
| Brackish water, without wheat | EM | 10<sup>4</sup> | 3 | 24.73 |
| Brackish water, with wheat | S | 10<sup>2</sup> | 1 | 38.23 |
| Brackish water, with wheat | S | 10<sup>2</sup> | 2 | 36.88 |
| Brackish water, with wheat | S | 10<sup>2</sup> | 3 | 39.09 |
| Brackish water, with wheat | PEG | 10<sup>2</sup> | 1 | 31.85 |
| Brackish water, with wheat | PEG | 10<sup>2</sup> | 2 | 29.34 |
| Brackish water, with wheat | PEG | 10<sup>2</sup> | 3 | 31.79 |
| Brackish water, with wheat | AC | 10<sup>2</sup> | 1 | 30.74 |
| Brackish water, with wheat | AC | 10<sup>2</sup> | 2 | 30.72 |
| Brackish water, with wheat | AC | 10<sup>2</sup> | 3 | 31.16 |
| Brackish water, with wheat | AC/PEG | 10<sup>2</sup> | 1 | 29.05 |
| Brackish water, with wheat | AC/PEG | 10<sup>2</sup> | 2 | 30.94 |
| Brackish water, with wheat | AC/PEG | 10<sup>2</sup> | 3 | 30.24 |
| Brackish water, with wheat | EM | 10<sup>2</sup> | 1 | 27.31 |
| Brackish water, with wheat | EM | 10<sup>2</sup> | 2 | 30.88 |
| Brackish water, with wheat | EM | 10<sup>2</sup> | 3 | 30.97 |
| Brackish water, with wheat | S | 10<sup>3</sup> | 1 | 33.23 |
| Brackish water, with wheat | S | 10<sup>3</sup> | 2 | 33.33 |
| Brackish water, with wheat | S | 10<sup>3</sup> | 3 | 33.70 |
| Brackish water, with wheat | PEG | 10<sup>3</sup> | 1 | 24.87 |
| Brackish water, with wheat | PEG | 10<sup>3</sup> | 2 | 32.53 |
| Brackish water, with wheat | PEG | 10<sup>3</sup> | 3 | 24.53 |
| Brackish water, with wheat | AC | 10<sup>3</sup> | 1 | 28.29 |
| Brackish water, with wheat | AC | 10<sup>3</sup> | 2 | 27.70 |
| Brackish water, with wheat | AC | 10<sup>3</sup> | 3 | 27.88 |
| Brackish water, with wheat | AC/PEG | 10<sup>3</sup> | 1 | 27.39 |
| Brackish water, with wheat | AC/PEG | 10<sup>3</sup> | 2 | 27.68 |
| Brackish water, with wheat | AC/PEG | 10<sup>3</sup> | 3 | 28.08 |
| Brackish water, with wheat | EM | 10<sup>3</sup> | 1 | 26.68 |
| Brackish water, with wheat | EM | 10<sup>3</sup> | 2 | 25.67 |
| Brackish water, with wheat | EM | 10<sup>3</sup> | 3 | 27.00 |
| Brackish water, with wheat | S | 10<sup>4</sup> | 1 | 29.55 |
| Brackish water, with wheat | S | 10<sup>4</sup> | 2 | 30.42 |
| Brackish water, with wheat | S | 10<sup>4</sup> | 3 | 31.34 |
| Brackish water, with wheat | PEG | 10<sup>4</sup> | 1 | 29.19 |
| Brackish water, with wheat | PEG | 10<sup>4</sup> | 2 | 26.10 |
| Brackish water, with wheat | PEG | 10<sup>4</sup> | 3 | 27.68 |
| Brackish water, with wheat | AC | 10<sup>4</sup> | 1 | 24.03 |
| Brackish water, with wheat | AC | 10<sup>4</sup> | 2 | 24.61 |
| Brackish water, with wheat | AC | 10<sup>4</sup> | 3 | 25.21 |
| Brackish water, with wheat | AC/PEG | 10<sup>4</sup> | 1 | 25.12 |
| Brackish water, with wheat | AC/PEG | 10<sup>4</sup> | 2 | 24.98 |
| Brackish water, with wheat | AC/PEG | 10<sup>4</sup> | 3 | 24.25 |
| Brackish water, with wheat | EM | 10<sup>4</sup> | 1 | 24.31 |
| Brackish water, with wheat | EM | 10<sup>4</sup> | 2 | 25.10 |
| Brackish water, with wheat | EM | 10<sup>4</sup> | 3 | 23.69 |

**Supplementary Table S9.**
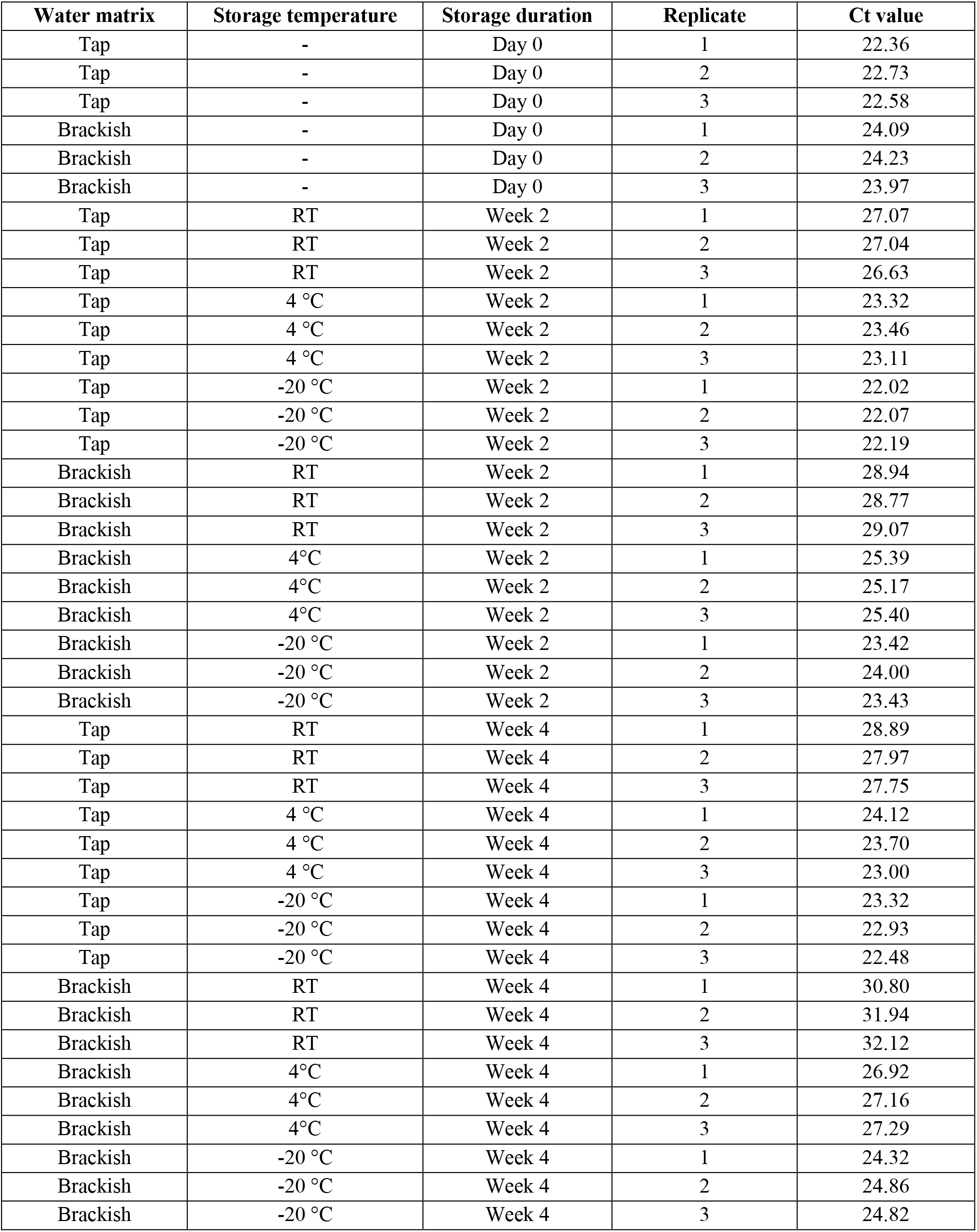
Individual Ct values obtained after enrichment of H4N6-spiked water samples using the AC/PEG method stored under three temperature conditions (room temperature, 4 °C, and −20 °C) for 0, 2, and 4 weeks in two water matrices (tap and brackish). Ct values are shown for each independent replicate. Abbreviations: RT, room temperature.

| **Water matrix** | **Storage temperature** | **Storage duration** | **Replicate** | **Ct value** |
|---|---|---|---|---|
| Tap | - | Day 0 | 1 | 22.36 |
| Tap | - | Day 0 | 2 | 22.73 |
| Tap | - | Day 0 | 3 | 22.58 |
| Brackish | - | Day 0 | 1 | 24.09 |
| Brackish | - | Day 0 | 2 | 24.23 |
| Brackish | - | Day 0 | 3 | 23.97 |
| Tap | RT | Week 2 | 1 | 27.07 |
| Tap | RT | Week 2 | 2 | 27.04 |
| Tap | RT | Week 2 | 3 | 26.63 |
| Tap | 4 °C | Week 2 | 1 | 23.32 |
| Tap | 4 °C | Week 2 | 2 | 23.46 |
| Tap | 4 °C | Week 2 | 3 | 23.11 |
| Tap | -20 °C | Week 2 | 1 | 22.02 |
| Tap | -20 °C | Week 2 | 2 | 22.07 |
| Tap | -20 °C | Week 2 | 3 | 22.19 |
| Brackish | RT | Week 2 | 1 | 28.94 |
| Brackish | RT | Week 2 | 2 | 28.77 |
| Brackish | RT | Week 2 | 3 | 29.07 |
| Brackish | 4°C | Week 2 | 1 | 25.39 |
| Brackish | 4°C | Week 2 | 2 | 25.17 |
| Brackish | 4°C | Week 2 | 3 | 25.40 |
| Brackish | -20 °C | Week 2 | 1 | 23.42 |
| Brackish | -20 °C | Week 2 | 2 | 24.00 |
| Brackish | -20 °C | Week 2 | 3 | 23.43 |
| Tap | RT | Week 4 | 1 | 28.89 |
| Tap | RT | Week 4 | 2 | 27.97 |
| Tap | RT | Week 4 | 3 | 27.75 |
| Tap | 4 °C | Week 4 | 1 | 24.12 |
| Tap | 4 °C | Week 4 | 2 | 23.70 |
| Tap | 4 °C | Week 4 | 3 | 23.00 |
| Tap | -20 °C | Week 4 | 1 | 23.32 |
| Tap | -20 °C | Week 4 | 2 | 22.93 |
| Tap | -20 °C | Week 4 | 3 | 22.48 |
| Brackish | RT | Week 4 | 1 | 30.80 |
| Brackish | RT | Week 4 | 2 | 31.94 |
| Brackish | RT | Week 4 | 3 | 32.12 |
| Brackish | 4°C | Week 4 | 1 | 26.92 |
| Brackish | 4°C | Week 4 | 2 | 27.16 |
| Brackish | 4°C | Week 4 | 3 | 27.29 |
| Brackish | -20 °C | Week 4 | 1 | 24.32 |
| Brackish | -20 °C | Week 4 | 2 | 24.86 |
| Brackish | -20 °C | Week 4 | 3 | 24.82 |

**Supplementary Table S10.** AIV screening results for the 14 brackish water samples screened before their use in the preparation of experimental samples. Values represent Ct values obtained from two independent measurements for each sample and method. Abbreviations: NA, not detected.

| Sample | S Ct | PEG Ct | AC Ct | AC/PEG Ct | EM Ct |
|---|---|---|---|---|---|
| 1 | NA / NA | NA / 37.5 | 39.5 / NA | 37.5 / NA | NA / NA |
| 2 | NA / NA | NA / 33.1 | 33.2 / NA | 34.3 / 36.7 | NA / NA |
| 3 | NA / NA | 32.3 / 32.0 | 35.8/ NA | NA / NA | NA / NA |
| 4 | 36.7 / NA | 36.5 / 35.1 | NA / NA | 31.5 / 32.2 | NA / NA |
| 5 | NA / NA | 29.3 / 36.4 | NA / NA | 35.1 / 34.8 | NA / NA |
| 6 | NA / NA | NA / NA | NA / NA | NA / NA | NA / NA |
| 7 | NA / NA | 35.8 / NA | NA / NA | NA / NA | NA / NA |
| 8 | NA / NA | NA / NA | NA / NA | NA / NA | NA / NA |
| 9 | NA / NA | NA / NA | 36.4 / 38.4 | 35.4 / 36.7 | Not tested |
| 10 | NA / NA | NA / NA | 35.6 / 37.5 | 35.4 / 35.6 | Not tested |
| 11 | NA / NA | NA / NA | 38.4 / NA | 35.9 / 36.4 | Not tested |
| 12 | Not tested | NA / NA | NA / NA | NA / NA | Not tested |
| 13 | Not tested | NA / NA | NA / NA | NA / NA | Not tested |
| 14 | Not tested | NA / NA | NA / NA | NA / 37.9 | Not tested |

## References

Abolmaaty A, Gu W, Witkowsky R, Levin RE. 2007. The use of activated charcoal for the removal of PCR inhibitors from oyster samples. J Microbiol Methods 68:349–352. 10.1016/j.mimet.2006.09.012

Ahrens AK, Selinka H, Wylezich C, Wonnemann H, Sindt O, Hellmer HH, Pfaff F, Höper D, Mettenleiter TC, Beer M, Harder TC. 2023. Investigating environmental matrices for use in avian influenza virus surveillance—surface water, sediments, and avian fecal samples. Microbiol Spectr 11:e02664–22. 10.1128/spectrum.02664-22

Azeem S, Guo B, Sato Y, Gauger PC, Wolc A, Yoon K-J. 2023. Utility of feathers for avian influenza virus detection in commercial poultry. Pathogens 12:1425. 10.3390/pathogens12121425

Borchardt MA, Boehm AB, Salit M, Spencer SK, Wigginton KR, Noble RT. 2021. The environmental microbiology minimum information (EMMI) guidelines: qPCR and dPCR quality and reporting for environmental microbiology. Environ Sci Technol 55:10210–10223. 10.1021/acs.est.1c01767

Cashdollar JL, Wymer L. 2013. Methods for primary concentration of viruses from water samples: a review and meta-analysis of recent studies. J Appl Microbiol 115:1–11. 10.1111/jam.12143

Crank K, Papp K, Barber C, Wang P, Bivins A, Gerrity D. 2023. Correspondence on “The Environmental Microbiology Minimum Information (EMMI) guidelines: qPCR and dPCR quality and reporting for environmental microbiology”. Environ Sci Technol 57:20448–20449. 10.1021/acs.est.3c07968

Coombe M, Iwasawa S, Byers KA, Prystajecky N, Hsiao W, Patrick DM, Himsworth CG. 2021. A systematic review and narrative synthesis of the use of environmental samples for the surveillance of avian influenza viruses in wild waterbirds. J Wildl Dis 57:1–18. 10.7589/JWD-D-20-00082

Deboosere N, Horm SV, Pinon A, Gachet J, Coldefy C, Buchy P, Vialette M. 2011. Development and validation of a concentration method for the detection of influenza A viruses from large volumes of surface water. Appl Environ Microbiol 77:3802–3808. 10.1128/AEM.02484-10

Farkas K, Pellett C, Alex-Sanders N, Bridgman MTP, Corbishley A, Grimsley JMS, Kasprzyk-Hordern B, Kevill JL, Pântea I, Richardson-O’Neill IS, Lambert-Slosarska K, Woodhall N, Jones DL. 2022. Comparative assessment of filtration- and precipitation-based methods for the concentration of SARS-CoV-2 and other viruses from wastewater. Microbiol Spectr 10:e01102–22. 10.1128/spectrum.01102-22

Fereidouni S, Starick E, Karamendin K, Di Genova C, Scott SD, Khan Y, Harder T, Kydyrmanov A. 2023. Genetic characterization of a new candidate hemagglutinin subtype of influenza A viruses. Emerg Microbes Infect 12:2225645. 10.1080/22221751.2023.2225645

Fox J, Weisberg S. 2019. An R Companion to Applied Regression, 3rd ed. Sage, Thousand Oaks, CA. https://www.john-fox.ca/Companion/

Germeraad EA, Elbers ARW, de Bruijn ND, Heutink R, van Voorst W, Hakze-van der Honing R, Bergervoet SA, Engelsma MY, van der Poel WHM, Beerens N. 2020. Detection of low pathogenic avian influenza virus subtype H10N7 in poultry and environmental water samples during a clinical outbreak in commercial free-range layers, Netherlands 2017. Front Vet Sci 7:237. 10.3389/fvets.2020.00237

Gouthro MT, Hayes EK, Prest T, Gagnon GA. 2025. Detection of avian influenza virus in surface waters using passive samplers. npj Viruses 3:57. 10.1038/s44298-025-00141-w

Hassan KE, Ahrens AK, Ali A, El-Kady MF, Hafez HM, Mettenleiter TC, Beer M, Harder T. 2022. Improved subtyping of avian influenza viruses using an RT-qPCR-based low density array: ‘Riems Influenza A Typing Array’, version 2 (RITA-2). Viruses 14:415. 10.3390/v14020415

Hénaux V, Samuel MD. 2011. Avian influenza shedding patterns in waterfowl: implications for surveillance, environmental transmission, and disease spread. J Wildl Dis 47:566–578. 10.7589/0090-3558-47.3.566

Hoffmann B, Depner K, Schirrmeier H, Beer M. 2006. A universal heterologous internal control system for duplex real-time RT-PCR assays used in a detection system for pestiviruses. J Virol Methods 136:200–209. 10.1016/j.jviromet.2006.05.020

Ho J, Stange C, Suhrborg R, Wurzbacher C, Drewes JE, Tiehm A. 2022. SARS-CoV-2 wastewater surveillance in Germany: long-term RT-digital droplet PCR monitoring, suitability of primer/probe combinations and biomarker stability. Water Res 210:117977. 10.1016/j.watres.2021.117977

Holzer C, Ho J, Tiehm A, Stange C. 2025. Wastewater monitoring—passive sampling for the detection of SARS-CoV-2 and antibiotic resistance genes in wastewater. Sci Total Environ 959:178244. 10.1016/j.scitotenv.2024.178244

Hokajärvi A-M, Rytkönen A, Tiwari A, Kauppinen A, Oikarinen S, Lehto KM, Kankaanpää A, Gunnar T, Al-Hello H, Blomqvist S, Miettinen IT, Savolainen-Kopra C, Pitkänen T. 2021. The detection and stability of the SARS-CoV-2 RNA biomarkers in wastewater influent in Helsinki, Finland. Sci Total Environ 770:145274. 10.1016/j.scitotenv.2021.145274

Hood G, Roche X, Brioudes A, von Dobschuetz S, Fasina FO, Kalpravidh W, Makonnen Y, Lubroth J, Sims L. 2021. A literature review of the use of environmental sampling in the surveillance of avian influenza viruses. Transbound Emerg Dis 68:110–126. 10.1111/tbed.13633

Hubbard LE, Givens CE, Stelzer EA, Killian ML, Kolpin DW, Szablewski CM, Poulson RL. 2023. Environmental surveillance and detection of infectious highly pathogenic avian influenza virus in Iowa wetlands. Environ Sci Technol Lett 10:1181–1187. 10.1021/acs.estlett.3c00668

Islam G, Gedge A, Lara-Jacobo L, Kirkwood A, Simmons D, Desaulniers J-P. 2022. Pasteurization, storage conditions and viral concentration methods influence RT-qPCR detection of SARS-CoV-2 RNA in wastewater. Sci Total Environ 821:153228. 10.1016/j.scitotenv.2022.153228

Juel MAI, Stark N, Nicolosi B, Lontai J, Lambirth K, Schlueter J, Gibas C, Munir M. 2021. Performance evaluation of virus concentration methods for implementing SARS-CoV-2 wastewater based epidemiology emphasizing quick data turnaround. Sci Total Environ 801:149656. 10.1016/j.scitotenv.2021.149656

Kaerber G. 1931. Beitrag zur kollektiven Behandlung pharmakologischer Reihenversuche. Arch Exp Pathol Pharmakol 162:480–483. 10.1007/BF01863914

Lenth R, Piaskowski J. 2025. emmeans: Estimated Marginal Means, aka Least-Squares Means. R package version 2.0.1. https://CRAN.R-project.org/package=emmeans

Munguía-Ramírez B, Armenta-Leyva B, Giménez-Lirola L, Zhang Y, Arruda B, Ciacci-Zanella G, Zimmerman J. 2026. Pilot assessment of RNA stabilization methods for influenza A virus in swine oral fluids. Pathogens 15:439. 10.3390/pathogens15040439

Munster VJ, Baas C, Lexmond P, Waldenström J, Wallensten A, Fransson T, Rimmelzwaan GF, Beyer WEP, Schutten M, Olsen B, Osterhaus ADME, Fouchier RAM. 2007. Spatial, temporal, and species variation in prevalence of influenza A viruses in wild migratory birds. PLoS Pathog 3:e61. 10.1371/journal.ppat.0030061

Okuya K, Kawabata T, Nagano K, Tsukiyama-Kohara K, Kusumoto I, Takase K, Ozawa M. 2015. Isolation and characterization of influenza A viruses from environmental water at an overwintering site of migratory birds in Japan. Arch Virol 160:3037–3052. 10.1007/s00705-015-2610-0

Pannwitz G, Wolf C, Harder T. 2009. Active surveillance for avian influenza virus infection in wild birds by analysis of avian fecal samples from the environment. J Wildl Dis 45:512–518. 10.7589/0090-3558-45.2.512

Panzarin V, Crimaudo M, Bonfante F, Marciano S, Berto P, Bofill-Mas S, Rusiñol M, Mazzetto E, Bortolami A, Fornasiero D, Martelli L, Mulatti P, Terregino C. 2025. Exploring the Use of Passive Samplers for the Surveillance of Avian Influenza Viruses in Wetlands: A Laboratory and Field Validation Study. Food Environ Virol 17:37. 10.1007/s12560-025-09649-z

Pedersen T. 2024. patchwork: The Composer of Plots. R package version 1.2.0. https://CRAN.R-project.org/package=patchwork

Pérez-Ramírez E, Acevedo P, Allepuz A, Gerrikagoitia X, Alba A, Busquets N, Díaz-Sánchez S, Álvarez V, Abad FX, Barral M, et al. 2012. Ecological factors driving avian influenza virus dynamics in Spanish wetland ecosystems. PLoS One 7:e46418. 10.1371/journal.pone.0046418

R Core Team. 2023. R: A language and environment for statistical computing. R Foundation for Statistical Computing, Vienna, Austria. https://www.R-project.org/.

Rondi FI, Neumann-Heise J, Harder T, Allendorf V, Depner K, Homeier-Bachmann T, Breuer L, Knauf S, Dreyer S, Globig A. 2026. Water-wheat-wait: monitoring wild mallards (*Anas platyrhynchos*) for avian influenza and ESBL-*E. coli* using feed-baited water bins, Germany, 2022 to 2023. One Health 23:101548. 10.1016/j.onehlt.2026.101548

Ryba S, Tacner J, Havlickova M, Stopka P. 2012. Stability of influenza virus as evaluated by integrity of its RNA. Acta Virol 56:125–128. 10.4149/av_2012_02_125

Schrader C, Schielke A, Ellerbroek L, Johne R. 2012. PCR inhibitors—occurrence, properties and removal. J Appl Microbiol 113:1014–1026. 10.1111/j.1365-2672.2012.05384.x

Spackman E, Senne DA, Myers TJ, Bulaga LL, Garber LP, Perdue ML, Lohman K, Daum LT, Suarez DL. 2002. Development of a real-time reverse transcriptase PCR assay for type A influenza virus and the avian H5 and H7 hemagglutinin subtypes. J Clin Microbiol 40:3256–3260. 10.1128/JCM.40.9.3256-3260.2002

Truyens CS, Berendes DM, Cantrell ME, Kossik AL, McCarthy KM, Mehrotra AS, Murphy JL, Pillay S, Raj SJ, Ramaswamy MS, Yakubu H, Holm RH. 2025. Advancing wastewater and environmental surveillance in LMICs for public health response and SDG data gaps. npj Clean Water 8:95. 10.1038/s41545-025-00523-w

Wickham H. 2016. ggplot2: Elegant Graphics for Data Analysis. Springer-Verlag, New York, NY. 10.1007/978-3-319-24277-4

Wickham H, Bryan J. 2023. readxl: Read Excel Files. R package version 1.4.3. https://CRAN.R-project.org/package=readxl

Wickham H, François R, Henry L, Müller K, Vaughan D. 2023. dplyr: A Grammar of Data Manipulation. R package version 1.1.4. https://CRAN.R-project.org/package=dplyr

Wickham H, Pedersen T, Seidel D. 2023. scales: Scale Functions for Visualization. R package version 1.3.0. https://CRAN.R-project.org/package=scales.

World Health Organization. 2011. Manual for the laboratory diagnosis and virological surveillance of influenza. Available from: https://www.who.int/publications/i/item/manual-for-the-laboratory-diagnosis-and-virological-surveillance-of-influenza

World Organisation for Animal Health. 2023. Terrestrial animal health code: chapter 10.4. Infection with high pathogenicity avian influenza viruses. Available from: https://www.woah.org/fileadmin/Home/eng/Health_standards/tahc/2023/chapitre_avian_influenza_viruses.pdf

